# High-resolution single-cell RNA sequencing using canFam4 reveals novel immune subsets and checkpoint programs in healthy dogs

**DOI:** 10.1101/2025.09.06.674668

**Authors:** Myung-Chul Kim, Taeeun Gu, Hyeewon Seo, Yewon Moon, Nicholas Borcherding, Ryan Kolb, Youngmin Yun, Woo-Jin Song, Chung-Young Lee, Hyun Je Kim, Weizhou Zhang

**Affiliations:** Veterinary Laboratory Medicine, Clinical Pathology, College of Veterinary Medicine, Jeju National University, Jeju 63243, Republic of Korea; Research Institute of Veterinary Medicine, College of Veterinary Medicine, Jeju National University, Jeju 63243, Republic of Korea; Department of Biomedical Sciences, Seoul National University Graduate School, Seoul 03080, Republic of Korea; Department of Pathology & Immunology, Washington University School of Medicine in St. Louis, St Louis, MO 63110, USA; Department of Pathology, Immunology and Laboratory Medicine, University of Florida College of Medicine, Gainesville, FL 32610, USA; Laboratory of Veterinary Internal Medicine, College of Veterinary Medicine, Jeju National University, Jeju 63243, Republic of Korea; Smart Animal Hospital, Seoul 06026, Republic of Korea; Department of Microbiology, School of Medicine, Kyungpook National University, Jung-gu, Daegu, Republic of Korea; Untreatable Infectious Disease Institute, Kyungpook National University, Daegu 41944, Republic of Korea; Genomic Medicine Institute, Seoul National University Medical Research Center, Seoul 03080, Republic of Korea; Department of Microbiology and Immunology, Seoul National University College of Medicine, Seoul 03080, Republic of Korea; UF Health Cancer Center, University of Florida, Gainesville, FL 32610, USA

**Keywords:** canFam4, circulating leukocytes, reference transcriptome, scRNA-seq, translational model

## Abstract

Single-cell RNA sequencing (scRNA-seq) enables high-resolution profiling of immune heterogeneity. Although previous studies have mapped the single-cell transcriptomic atlases of peripheral leukocytes in healthy dogs, the identification and functional characterization of distinct immune subsets remain incomplete. We constructed a single-cell atlas of peripheral leukocytes from six healthy small-breed dogs using the 10x Genomics platform and the updated canFam4 genome. Analysis of 30,040 high-quality transcriptomes revealed 51 distinct immune subsets, including *CD14*□*CD33*□ monocytes, *XCR1*□*CD1D*□ dendritic cells, *CEACAM1*□*CD24*□ neutrophils, and *IL32*□*BATF*□ regulatory T cells, which were underrepresented in canFam3.1-based studies. Interferon-enriched *CD14*□ monocytes and *CD4*□ T subsets associated with myxomatous mitral valve disease were also identified. Functional analysis revealed that *PDCD1* attenuates TCR signaling, *LAG3* modulates malate metabolism in *CD4*□ T cells, and suppresses *TBX21* in *CD8*□ T cells associated with viral response. *CD274* encoding PD-L1 was linked to IL-10 production in neutrophils, and *CTLA4* represented an initial activation of double-negative T subsets. T cell exhaustion scores and proliferative fractions varied across cohorts, reflecting differences in environmental antigenic exposures. Our study represents the first comprehensive, gene-resolved single-cell analysis that reveals immunoregulatory checkpoint mechanisms underlying immune homeostasis in healthy dogs. Our dataset will serve as a valuable resource for future comparative and translational immunology research in dogs.

## Introduction

Humans and dogs inextricably share not only genetic traits but also environments, lifestyles, stressors, and microbial exposures (1,2). Moreover, dogs have many of the same naturally occurring diseases as humans (3), including as cancer, inflammatory bowel disease, and cardiomyopathy (2,4). Consequently, companion dogs serve as valuable spontaneous models for translating basic science into clinical applications (5). Cross-fertilization between veterinary and human medicine is most advanced in oncology (6).

Single-cell RNA sequencing (scRNA-seq) dissects dynamic and heterogeneous tissue microenvironments by characterizing the transcriptome at the single-cell level, thereby improving our understanding of cell identity, fate, and function in the context of both normal biology and pathology (7). To date, scRNA-seq has begun to unlock the secrets of veterinary diseases with application to canine cells derived from blood (8–13), lymphoid tissues (14–16), bronchoalveolar lavage (17), hippocampus (18), liver (16), lung (19), adipose tissues (20), inflamed tissues (12,21–23), and cancers (10,24–27). More recently, the single-cell transcriptome atlas has elucidated canine hematopoiesis (28) and peri-implantitis pathogenesis via pseudotime and interactome analyses (29). Among various sample origin, peripheral blood mononuclear cells (PBMC) have served as a easily accessible and valuable source for gaining insight into the tumor microenvironment (8,13), lymphocyte clonal expansion (10), inflammatory disease etiology (23), innate immunity (16), aging (20), and preclinical immunotherapy optimization (12).

Although the canine genome is fully sequenced and annotated, the widely used reference or canFam3.1 remains incomplete with reads unmapped (30), lacking some key cell lineage and functional genes, such as *CD14*, *FCGR3A*, and the CD1 family of genes (8–12). Recently, new canine genome references, such as GSD_1.0 (canFam4), have been released, offering potential for improved transcriptional resolution with high contiguity (30). Evaluating canFam4 in the context of scRNA-seq analysis could have the potential to enhance single-cell recovery and cell-type identification, which have not been reported yet.

Prior studies have profiled canine leukocytes using scRNA-seq (8–10,23). However, the identification and functional characterization of distinct immune subsets remain incomplete in dogs, and it remains unclear whether all clinically relevant subsets have been fully uncovered. To address these gaps, we performed scRNA-seq on PBMCs from clinicopathologically healthy, client-owned, indoor small-breed dogs. For the first time, we demonstrate that using canFam4 significantly increases cell recovery and reveals previously undetected markers, including *CD14*. In addition, we provide immune subsets potentially involved in disease and regulatory mechanisms of immune checkpoint genes, such as *PDCD1*, *CTLA4*, *LAG3*, *CTLA4*, and *CD274* in dogs. To our knowledge, this is the first scRNA-seq study to provide molecular evidence of how canine immune subsets maintain homeostasis. Our dataset and methodology will serve as foundational resources for future investigations into canine inflammatory diseases, including cancers and autoimmunity.

## Materials and Methods

### Study subject enrollment and inclusion criteria

Client-owned adult dogs (7 to 12 years old) of common domestic breeds were enrolled from the Veterinary Medical Teaching Hospital at Jeju National University (Jeju-Si, South Korea). All dogs were housed indoors and confirmed by their owners to have no preexisting disease conditions. Inclusion criteria included no clinical signs of disease on physical examination, normal clinicopathologic results, and no vaccinations or treatments within four weeks before blood sampling. Written informed consent was obtained from each owner before enrollment. The study protocol was reviewed and approved by the Jeju National University Institutional Animal Care and Use Committee (IACUC No. 20230072).

### Clinicopathologic examinations

Resident veterinarians conducted thorough physical examinations and clinical assessments. Hematological and biochemical analyses were performed using the ProCyte Dx and Catalyst One systems (IDEXX Laboratories, MA, USA), respectively. To screen for Babesia infection, we employed both a point-of-care antibody rapid test (Canine Babesia Antibody Rapid Kit, BioNote Inc., Gyeonggi-do, South Korea) and real-time PCR (CareDx™ Canine Babesia Real-Time PCR Kit, Carevet Inc., Gyeonggi-do, South Korea). Additionally, Anigen Rapid CaniV-4 (BioNote Inc.) and SNAP 4Dx Plus (IDEXX Laboratories) assays were used to detect other protozoal pathogens. Only dogs testing negative for all infectious agents were included in the study.

### Isolation of peripheral blood mononuclear cells

Blood was collected via jugular venipuncture and processed immediately. PBMCs were isolated by density gradient centrifugation using SepMate-15 tubes (Stemcell Technologies, Vancouver, Canada) and Ficoll-Paque□PLUS (GE Healthcare, Chicago, IL, USA). The PBMC fraction was washed twice with Dulbecco’s phosphate-buffered saline (DPBS; Thermo Fisher Scientific, Waltham, MA, USA) containing 2% heat-inactivated fetal bovine serum (FBS; Life Technologies, Pleasanton, CA, USA). A small aliquot of the suspension was cytospun onto glass slides and stained with Diff-Quik to confirm the purity and composition of isolated cells. The remaining cells were cryopreserved in a cytoprotective medium (Cellbanker□1, Zenogen Pharma, Koriyama, Japan) and stored in liquid nitrogen until further analysis.

### Fluorescence-activated cell sorting

Thawed PBMCs were washed and incubated on ice for 10□minutes with an anti-dog Fc receptor blocking reagent (Invitrogen, eBioscience, San Diego, CA, USA). After washing with ice-cold DPBS containing 2% FBS, cells were stained for 30□minutes at 37□°C, protected from light, with Fixable Viability Dye eFluor□780 (Invitrogen, eBioscience), anti-dog CD45 PE (clone YKIX□716.13; Bio-Rad Laboratories, Hercules, CA, USA), and anti-dog CD3 FITC (clone CA17.2A12; Bio-Rad Laboratories). Live single cells were then sorted on a FACSAria□III (BD Biosciences Pharmingen, San Diego, CA, USA), gating on CD45□CD3□ T cells and CD45□CD3□ non-T leukocytes. Post-sort viability and cell counts were assessed using a LUNA-FL™ Automated Fluorescence Cell Counter (Logos Biosystems, Anyang, South Korea). Equal numbers of CD45□CD3□ and CD45□CD3□ cells from each dog were pooled and immediately processed for single-cell library construction on the 10x Genomics Chromium platform in a single run.

### 10X Genomics single-cell RNA sequencing library construction

Single-cell libraries were prepared using the Chromium Next GEM Single Cell 5′ Kit v2 (10x Genomics) following the manufacturer’s protocol (31,32). Libraries were sequenced on an Illumina NovaSeq□6000 with 2□×□150□bp paired-end reads. Base calling and FASTQ generation were performed with bcl2fastq (Illumina) and cellranger mkfastq (Cell Ranger v8.0.0, 10x Genomics). Reads were aligned and quantified using cellranger count by using two canine genome references: CanFam3.1 (GCA_000002285.3) and GSD_1.0/CanFam4 (GCF_011100685.1). Genome references were constructed with cellranger mkref using filtered FASTA and GTF files containing only protein-coding genes.

### scRNA-seq data integration, initial preprocessing, and sub-clustering

All single-cell data were processed and analyzed in Seurat v5.1.0 (R v4.2). After loading each sample, low-quality cells were filtered out if they met any of the following criteria: fewer than 200 or more than 4,000 unique features and 10% mitochondrial gene content. Filtered datasets were normalized using Seurat’s default workflow. Next, all six samples were merged and integrated into a single Seurat object using 3,000 anchor features. Principal component analysis (PCA) was performed for dimensionality reduction, and principal components (PCs) 1–30, which were selected based on JackStraw significance, P value < 1e-5, and percent variance explained, were used for downstream clustering. We applied graph-based clustering on UMAP and t-SNE embeddings at a resolution of 3.7. To determine cohort-specific phenotypes, we incorporated publicly available canine scRNA-seq datasets, including peripheral blood TCR αβ T cells (GSE218355) (9) and PBMCs (GSE225599) (8). FASTQ files from GSE144730 – canine PBMCs in atopic dermatitis (23) – were not included due to unsupported library chemistry. Available external datasets were processed and integrated alongside our own using the same quality control, normalization, and anchor-based integration pipeline. Harmony integration (v1.2.0) was performed on the PCA-reduced data to correct batch effects across studies. The first 30 Harmony dimensions were subsequently used for neighbor finding, clustering, UMAP, and t-SNE visualization. Doublets were detected and removed with scDblFinder v1.18.0 (31). Single-cell clusters were annotated by canonical lineage markers, published signatures for rare populations, and unbiased cell-type inference via SingleR v2.6.0 (33) using celldex v1.14.0 reference panels. Finally, the escape v2.0.0 package (34) was used to calculate enrichment scores for specific immune gene signatures drawn from original canine studies (8,22).

### Differentially expressed gene analysis

Differential expression was assessed using Seurat’s likelihood-ratio test, comparing each cluster to all other cells. Marker genes for individual clusters were identified with FindAllMarkers, applying cutoffs of absolute log□-fold change >□0.25 and ≥□25% of cells in the cluster expressing the gene. To compare cluster markers and DEGs across experimental groups, we used FindMarkers with a more stringent threshold (absolute log□-fold change >□0.5 and *P*L<□0.05).

### Gene set enrichment analysis and Gene Ontology analysis

We performed gene set enrichment analysis (GSEA) using the escape R package, employing Hallmark gene sets from the Molecular Signatures Database and those from previous canine studies (35–39). Differentially expressed genes (DEGs) were further analyzed for Gene Ontology (GO) enrichment using both the PANTHER annotation (for *Canis lupus familiaris*) and the ShinyGO v0.80 web tool with species set to dog. GO terms were considered significant at *P*□<□0.05 and false discovery rate (FDR)□<□0.05. Pathway and gene set visualizations were generated with DittoSeq v1.4.4 and pheatmap v1.0.12.

### Pseudotime analysis

Pseudotime analysis was performed using Slingshot v2.12.0 based on UMAP embeddings and Seurat-derived clusters, as previously described (34). Naïve T cells were specified as the trajectory root. Multiple lineages were inferred, and gene expression dynamics were analyzed along lineage-specific pseudotime axes.

### Cell cycle analysis

Cell cycle phase was assigned using Seurat’s CellCycleScoring function with the cc.genes.updated.2019 gene set, following established protocols (31,32).

### Cell-to-cell interaction analysis

Intercellular communication was inferred from scRNA-seq data using the CellChat R package v2.1.2 (40). Group-specific CellChat objects were merged into a single master object for comparative analysis. Overall signaling networks were visualized with the netVisual_heatmap function. To identify and display significant ligand–receptor pairs, we applied the subsetCommunication and netVisual*_*bubble functions using thresholds of ligand log□-fold change >□0.2, receptor log□-fold change >□0.1, and *P*□<□0.01.

### Statistical methods and reproducibility

Statistical tests were performed primarily within Seurat v5.1.0. For single-cell differential expression, the nonparametric Wilcoxon rank-sum test was applied via FindAllMarkers and FindMarkers. Count-level mRNA comparisons employed one-way ANOVA and two-tailed unpaired Student’s *t*-tests (Welch’s correction where appropriate) using the ggpubr v0.6.0 package. To evaluate the correlation between two gene signature enrichment, both Pearson’s correlation and simple linear regression analyses were performed using R. Pearson’s correlation coefficient (r) and its statistical significance were computed using the cor.test() function. A simple linear regression analysis was performed using the lm() function. Results were considered statistically significant at *P*□<□0.05.

### Data and code availability

The raw and processed scRNA-seq data generated in this study have been deposited in the NCBI Gene Expression Omnibus (GEO) under accession number GSE301630. Custom analysis scripts and pipelines are available from the corresponding author upon reasonable request.

## Results

### Clinical and laboratory evaluation of the health status of the dogs

A summary of the demographic characteristics is provided in **Supplementary Data 1**. In brief, six healthy dogs (median age, 9.3 years) representing three common domestic breeds, including Poodle (n = 1), Maltese (n = 3), and mixed-breed (n = 2), were enrolled in this study. History taking, physical examination, and clinicopathologic analyses revealed no abnormalities (**Supplementary Data 2**). All subjects tested negative for infectious agents on a point-of-care immunoassay. Furthermore, resident veterinarians confirmed the absence of any clinical signs or disease during a six-month follow-up conducted via telephone.

### The study workflow and the quality control of the data

The study scheme (**Figure 1A**), library statistics (**Supplementary Data 3**), and standard preprocessing and quality control of the scRNA-seq data (**Supplementary Figures 1A, 1B, 1C**) are available. Using the 10x Genomics platform, a total of 30,040 cells were obtained from healthy 1 (n=5,659), healthy 2 (n=2,674), healthy 3 (n=4,833), healthy 4 (n=7,644), healthy 5 (n=6,384), and healthy 6 (n=2,846) groups (**Supplementary Figure 1D**). Immune cells across breed, age, and sex shared a similar global structure of RNA expression as revealed by dimensional reduction and clustering analyses (**Supplementary Figure 1E**). There were approximately 15.6% of potential doublets detected by scDblFinder (**Supplementary Figure 1F**), which turned out to be myeloid, T, and B lymphocyte complexes and were therefore excluded to ensure the integrity of downstream analyses (**Supplementary Figure 1G**).

**Figure 1.**
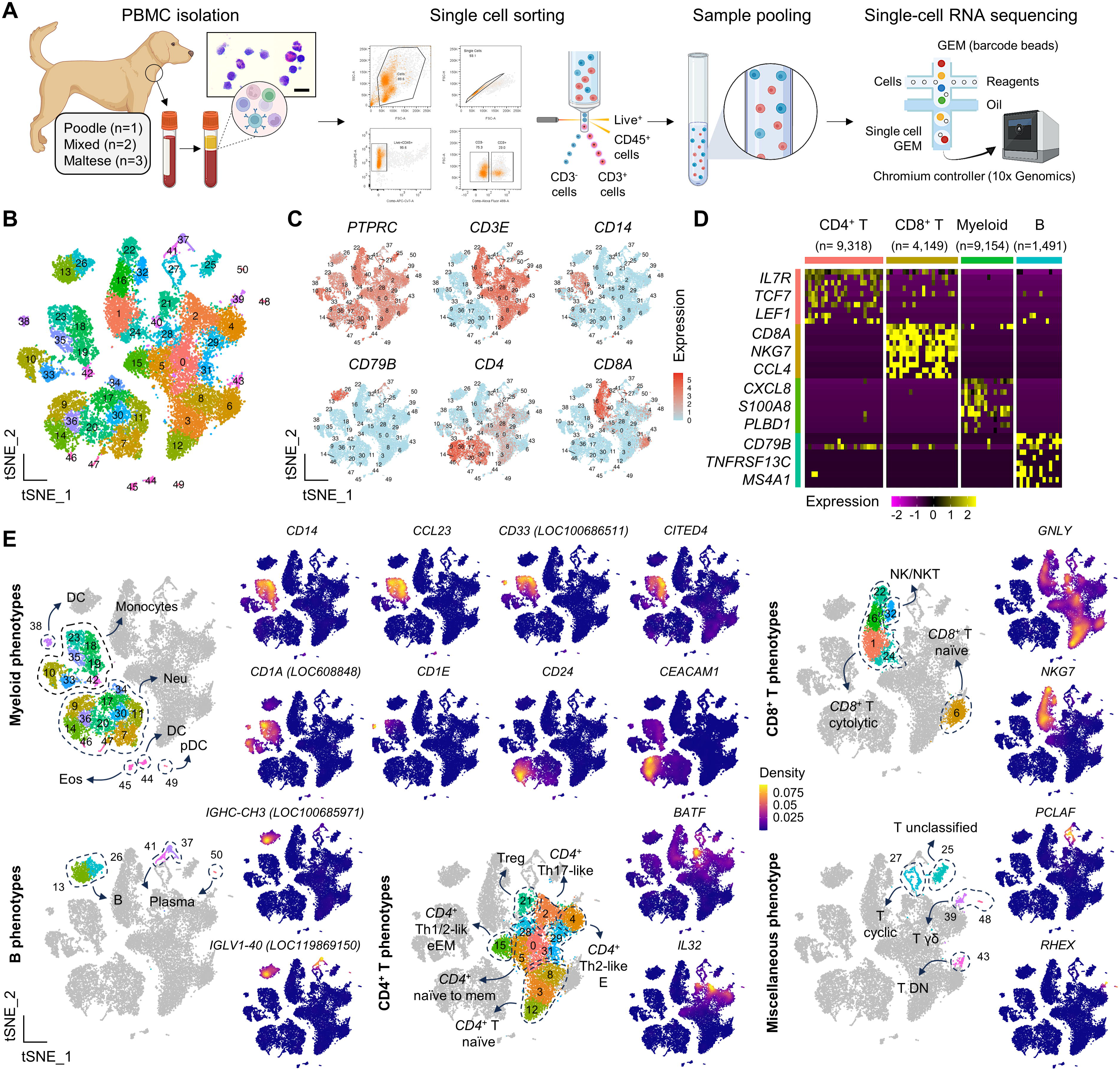
Study design and identification of major immune cell clusters by integrated scRNA-seq analysis in dogs. (A) Schematic overview of the study. Live, single CD3^+^ and CD3^-^ peripheral immune cells were isolated from the jugular vein of 6 clinically healthy dogs, flow-sorted, pooled, and subject to single-cell library preparation and downstream bioinformatic analysis. A representative photomicrograph of a hematoxylin and eosin-stained PBMC smear is shown at 400× magnification (scale bar = 20□μm). Scientific illustrations were created using BioRender under an academic license. (B) A total of 51 transcriptionally distinct clusters were identified in the integrated Seurat object and visualized on a tSNE plot. (C) Canonical lineage markers used to classify major immune cell types are shown on the tSNE feature plot. (D) Representative genes among the top 10 DEGs for each major lineage are visualized on a heatmap. The number of cells analyzed per lineage is indicated below. (E) tSNE density plots showing representative canonical markers used to identify functionally distinct immune subsets. All marker genes, except *FOXP3*, were annotated only in the canFam4 genome reference.

### Identification of major immune subsets in circulating leukocytes of healthy dogs

After singlet selection, we identified 25,355 cells across 51 clusters in the integrated Seurat object (**Figure 1B**). All immune subsets were assigned to each biological replicate (**Supplementary Figure 1H**). Clustering performance was evaluated by identifying the major immune cell types, showing a clear separation of T, B, and myeloid populations (**Figure 1C**). Out of the top 10 DEGs to define each major immune subset, representative genes were presented on the heatmap (**Figure 1D**). Representative genes defining functionally distinct immune subsets were identified (**Figure 1E, Supplementary Figure 2**). Unbiased cell type recognition, performed by enrichment analysis using gene signatures defining canine and human immune subsets, corroborated our cell type annotation (**Supplementary Figures 3A and 3B**). Overall, cell type and subset identification in this study supported previous scRNA-seq results applied to circulating leukocytes of healthy dogs. (8–10,23). For example, regulatory T cells (Treg; cluster 21) expressed high levels of *IL2RA* and *FOXP3*, along with a low level of *IL7R*. Plasma cells (clusters 37, 41, and 50) expressed *MZB1* and *JCHAIN*. Gamma delta (γδ) T cells (clusters 39 and 48) showed specific expression of *RHEX* and *SCART1* (*LOC491694*). Plasmacytoid dendritic cells (pDC; cluster 49) were defined by expression of *IL3RA* and *TCF4*. Myeloid cells expressed *DPYD*. Naïve T cells (clusters 3, 8, and 12) expressed *CCR7* and *SELL*. Activated T cells upregulated genes associated with co-receptors (*CD28* and *ICOS*), cytokines (*IL2* and *TNF*), and cytotoxic granules (*PRF1*, *GZMB*, and *GZMK*). Cycling T cells (cluster 27) expressed *MKI67* and *PCLAF*, with a high proportion of cells in S and G2/M phase (**Supplementary Figure 4**).

It is worth noting that adopting canFam4 enables the discovery of features not reported in previous scRNA-seq studies in dogs (8–10,23). **Figure 1E** presents representative functional clustering and lineage markers annotated in CanFam4. For example, *CD14*, *CCL23*, *CD33* (*LOC100686511*), *CD1E*, *CD1A*, *CD24*, *CITED4*, and *CEACAM1* were identified in monocytes, dendritic cells (DC), and neutrophils. Lymphocytic lineage and functional markers, such as *LOC607937* (T cell receptor alpha variable 9), *CD8B*, *GLNY NKG7*, *GZMH* (*LOC490629*), *BATF*, *IL32*, *LOC100685971* (immunoglobulin heavy chain constant region CH3), and *LOC119869150* (immunoglobulin lambda-1 light chain-like) were also identified.

We then conducted a more comprehensive analysis to investigate reference-specific highly variable features across genome builds. Using the same preprocessing, quality control, and integration pipelines, we applied canFam3.1 to generate a new master Seurat object. Interestingly, we identified 1,490 highly variable features specific to canFam4 (**Supplementary Data 4**), which were expressed across diverse immune subsets (**Supplementary Figure 5A**). In addition to revealing novel genes, canFam4 also improved scRNA-seq quality control metrics (**Supplementary Figure 5B**). When applied to the Seurat object, canFam4 significantly increased the number of assigned cells and decreased the proportion of non-human homologous genes. The number of RNA features and the proportion of mitochondrial genes remained consistent across references. In summary, we generated scRNA-seq libraries representing functionally distinct immune subsets in healthy dogs and improved transcriptional resolution and single-cell assignment accuracy by utilizing the canFam4 reference.

### Identification and characterization of myeloid subsets

Contrary to humans, granulocytes, polymorphonuclear cells (PMNs), or myeloid-derived suppressive cells (MDSCs) are collected during gradient-based isolation of canine peripheral blood (8,10,41). Canine myeloid population is highly heterogeneous in its transcriptional phenotypes, but it had not been fully understood at the single-cell level (8–10,23). Therefore, we first focused on identifying and characterizing functionally distinct myeloid subsets. Sub-clustering of *DPYD*□ myeloid cells revealed 23 clusters in dogs (**Figure 2A**).

**Figure 2.**
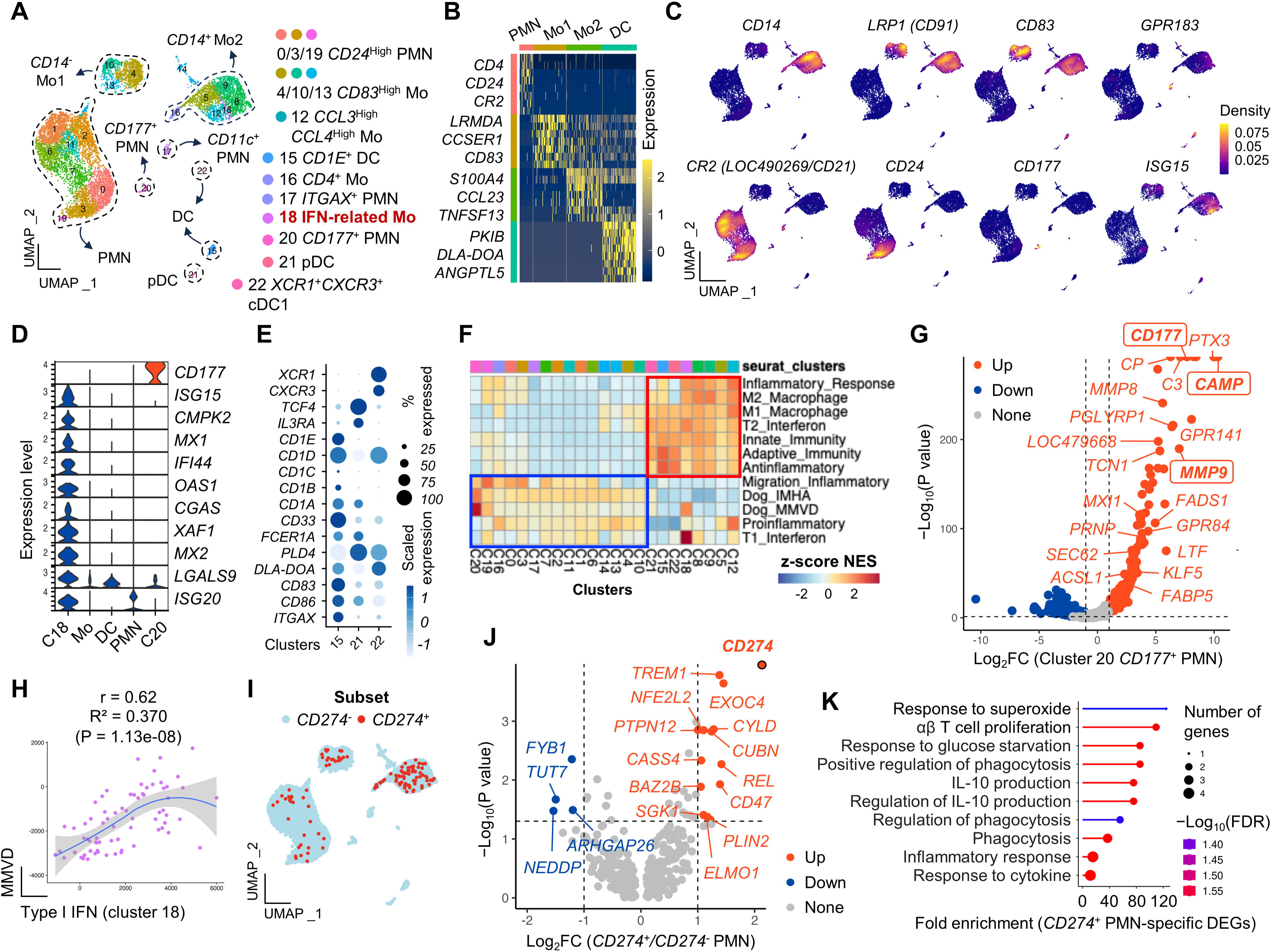
Identification and characterization of myeloid subpopulations by scRNA-seq. (A) UMAP visualization of 23 myeloid subsets classified into classical (*CD14*□) and non-classical (*CD14*□) monocytes, granulocytes (PMNs), and dendritic cells (DCs). Representative subsets are labeled. (B) Heatmap showing representative genes among the top 7 DEGs defining each cell type. (C) Density plots of selected marker genes distinguishing functionally distinct myeloid subsets. (D) Violin plots highlighting clusters 18 and 20, characterized by upregulation of IFN-related genes and *CD177*, respectively. (E) Dot plot showing canonical markers for functional dendritic cell subsets. (F) GSEA. Immune-related pathways are enriched in DC/classical monocyte clusters (red box; e.g., M1/M2 macrophage, IFN signatures) and in PMN/non-classical monocyte clusters (blue box; e.g., migration, Dog_IMHA, Dog_MMVD signatures). (G) Volcano plot showing DEGs in *CD177*□ PMNs compared to other PMN subsets. Genes associated with Dog_MMVD are highlighted. (H) Scatter plot showing the correlation between type I IFN and MMVD gene signature scores in cluster 18. (I) Feature plot displaying the distribution of *CD274*□ PMNs and monocytes. (J) Volcano plot showing representative DEGs in *CD274*□ PMNs compared to *CD274*□ PMNs. *CD274* is highlighted with a log□FC of 16.4 and P = 0. (K) GO analysis of *CD274*□ PMN-specific DEGs reveals enrichment in biological processes associated with T cell proliferation, glucose starvation, phagocytosis, and IL-10 production. P-values were derived using one-way analysis of variance implemented in the ggpubr R package and Wilcoxon rank-sum tests for pairwise comparisons. P < 0.05, P < 0.01, P < 0.001. Abbreviations: PMN, polymorphonuclear cells; FC, fold change; DEG, differentially expressed genes; C, cluster; Mo, monocytes; DC, dendritic cells; pDC, plasmacytoid DC; GO, gene ontology; GSEA, gene set enrichment analysis.

In this study, canine myeloid populations primarily consisted of granulocytes, monocytes, and DC subsets, exhibiting a distinct global RNA expression structure distinguished by canonical marker genes such as *CD4*, *CD24*, *CR2* (*LOC490269*/*CD21*), *S100A4*, *CD83*, *CCL23*, *TNFSF13*, and *DLA*-*DOA* (**Figure 2B**). Canonical and cluster-specific markers used to define functionally distinct subsets are shown in **Supplementary Figure 6A and Data 5**. Notably, we identified *S100A4*□*CCL23*□*TNFSF13*□ monocytes (clusters 5, 8, 9, 12, 16, and 18) that expressed *CD14* (**Figure 2C**), a canonical marker for classical monocytes which had not been identified in previous scRNA-seq studies (8–10,23). Clusters 4, 10, and 13 lacked *CD14* expression but expressed *CD83* and *CD91* (*LRP1*), suggesting potential non-classical monocytes (12,42). However, the *CD16* transcript was found to be unannotated in canFam4. Cluster 18 was further characterized by the expression of interferon (IFN)-stimulated genes, including *ISG15*, *MX1*, *MX2*, *IFI44*, *OAS1*, and *XAF1*, indicating IFN-related monocytes (31) (**Figure 2D**). A majority of the IFN-related monocytes showed *LGALS9* expression (**Supplementary Figure 6B**).

DC subsets (clusters 15, 21, and 22) specifically expressed *GPR183* alongside *DLA*-*DOA*, *CD86*, and *PLD4* (**Figures 2C, 2E, and Supplementary Figure 6A**). Importantly, all *CD1* family genes (*CD1A*, *CD1B*, *CD1C*, *CD1D*, and *CD1E*) were differentially expressed among the DC subsets (**Figure 2D**). Similar to humans, dogs exhibited *XCR1*□ terminally differentiated conventional type 1 DCs (cDC1; cluster 22) (43), *TCF4*□*IL3RA*□ pDC (cluster 21), and *ITGAX*□*FCER1A*□ cDC2 (cluster 15). The *XCR1*□ DC was characterized by significant *MIF* upregulation (**Supplementary Figure 6C**).

PMNs exclusively expressed *CD21* (*CR2***/***LOC490269*) and *CD4*, while *CD177* (cluster 20), *ITGAX* (cluster 17), *CEACAM1*, and *CD24* (clusters 0, 3, and 19) further delineated neutrophil subsets (**Figures 2C, 2E, and Supplementary Figure 6D**). Cluster 14 was likely a doublet, as it co-expressed both *CD14* and *CD3E*.

Next, we functionally characterized how these distinct myeloid subsets contribute to immune homeostasis in healthy dogs (**Figure 2F**). First, *CD14*□ monocytes and DC subsets were preferentially enriched in inflammatory gene signatures, such as M1/M2 macrophage markers, innate/adaptive immunity, and IFN signaling pathways (**Figure 2F, red box**). In contrast, neutrophil subsets were preferentially enriched for gene sets related to leukocyte migration involved in the inflammatory response (**Figure 2F, blue box**). We also found that certain myeloid subsets may be associated with immune dysregulation of canine diseases. For instance, *CD177*□ neutrophils showed strong enrichment in gene signatures associated with immune-mediated hemolytic anemia (IMHA) (35) and myxomatous mitral valvular disease (MMVD) (44). In the *CD177*□ neutrophils, there was also a significant positive correlation between IMHA and MMVD signatures (**Supplementary Figure 6E**). Genes such as *CD177*, *CAMP*, and *MMP9* were significantly upregulated in these neutrophils (**Figure 2G**). IFN-related monocytes also exhibited MMVD signature enrichment, with notable upregulation of *ISG15* and *MX1* (**Supplementary Figure 6F**). Additionally, there was a moderate strong significant positive correlation between type I IFN and MMVD gene signatures (**Figure 2H**).

Anti-PD-L1 antibody has antitumor effects in dogs with naturally occurring cancers (45,46), but their function in dogs remains poorly characterized. We investigated the contribution of *PD-L1* (encodes *CD274* protein) in myeloid-mediated immune regulation. At steady-state, *CD274* was expressed in neutrophils and monocytes but not in DCs (**Figure 2I and Supplementary Figure 6G**). We separated myeloid cells based on *CD274* expression and performed differential gene expression analysis. *CD274*□ neutrophils exhibited 18 differentially expressed genes compared to *CD274*□ neutrophils (**Figure 2J and Supplementary Data 6**). Gene ontology (GO) analysis of these DEGs revealed enrichment in pathways such as αβ T cell proliferation, IL-10 production, phagocytosis, and the inflammatory response, supporting the functional phenotype of PMN-MDSC (47) (**Figure 2K and Supplementary Data 7**). Notably, *CD47*, a part of the IL-10 signature, was significantly upregulated in *CD274*□ neutrophils. Similarly, 172 DEGs defined *CD274*□ monocytes (**Supplementary Figure 6G**), which were enriched in PD-L1 checkpoint, toll-like receptor, NF-κB, and C-type lectin receptor pathways (**Supplementary Figure 6I and Data 6, 7**). Taken together, we identified functionally distinct myeloid subsets, demonstrating their potential clinical relevance in regulating immune homeostasis in dogs.

### Identification and characterization of *CD4*^+^ T subsets

T helper subsets play major roles in canine health and disease (48). We next identified and characterized *CD4*□ T cell subsets in healthy dogs and evaluated their potential clinical relevance (**Figure 3A**). Representative marker genes used for annotation are listed in **Supplementary Figure 7A and Data 8**. Briefly, naïve T cells (clusters 0, 1, 11, 13, and 15) and effector**/**memory T cells (clusters 2, 3, 4, 5, 6, 7, 8, 12, 14, and 17) were distinguished based on *SELL* and *CD44* expression, canonical markers of naïve and memory phenotypes (48) (**Figure 3B**). Using canine *CD4*^+^ helper T signatures (8,22,24), we identified Th1 (cluster 3), Th2 (cluster 6), and Th17 (cluster 5) subsets (**Figure 3C**). These subsets exhibited some degree of transcriptional overlap (**Figure 3C**). Similar to myeloid cells, *CD4*□ T cells included an IFN-related subset (cluster 14), characterized by strong enrichment and expression of IFN-stimulated genes, including *XAF1*, *OAS1*, *MX1*, and *ISG15* (**Figures 3C, 3D, upper panel**). Cluster 12 showed high expression and density of *PDCD1* (encodes PD-1 protein) (**Figure 3E**). Tregs (clusters 7 and 17) expressed *FOXP3*, *IL2RA*, *CCR4*, and *CTLA4*, along with less well-characterized genes, such as *BATF* and *IL32* (**Supplementary Figures 7A and 7B**), which have not been previously reported in dogs. Meanwhile, *CTLA4* was broadly expressed across multiple *CD4*□ T subsets (**Figure 3E**). Cluster 9 expressed *CD8A*, suggesting a population of double-positive (DP) T cells. Clusters 8, 11, and 15 remained less well characterized, but showed high expression of *CCDC3*, *LOC119876465* (a putative RNA polymerase B transcription factor), and *STAT3*, respectively. Cluster 10 showed high expression of myeloid markers such as *CSF3R*, *DPYD*, and *S100A8*, suggesting potential doublets.

**Figure 3.**
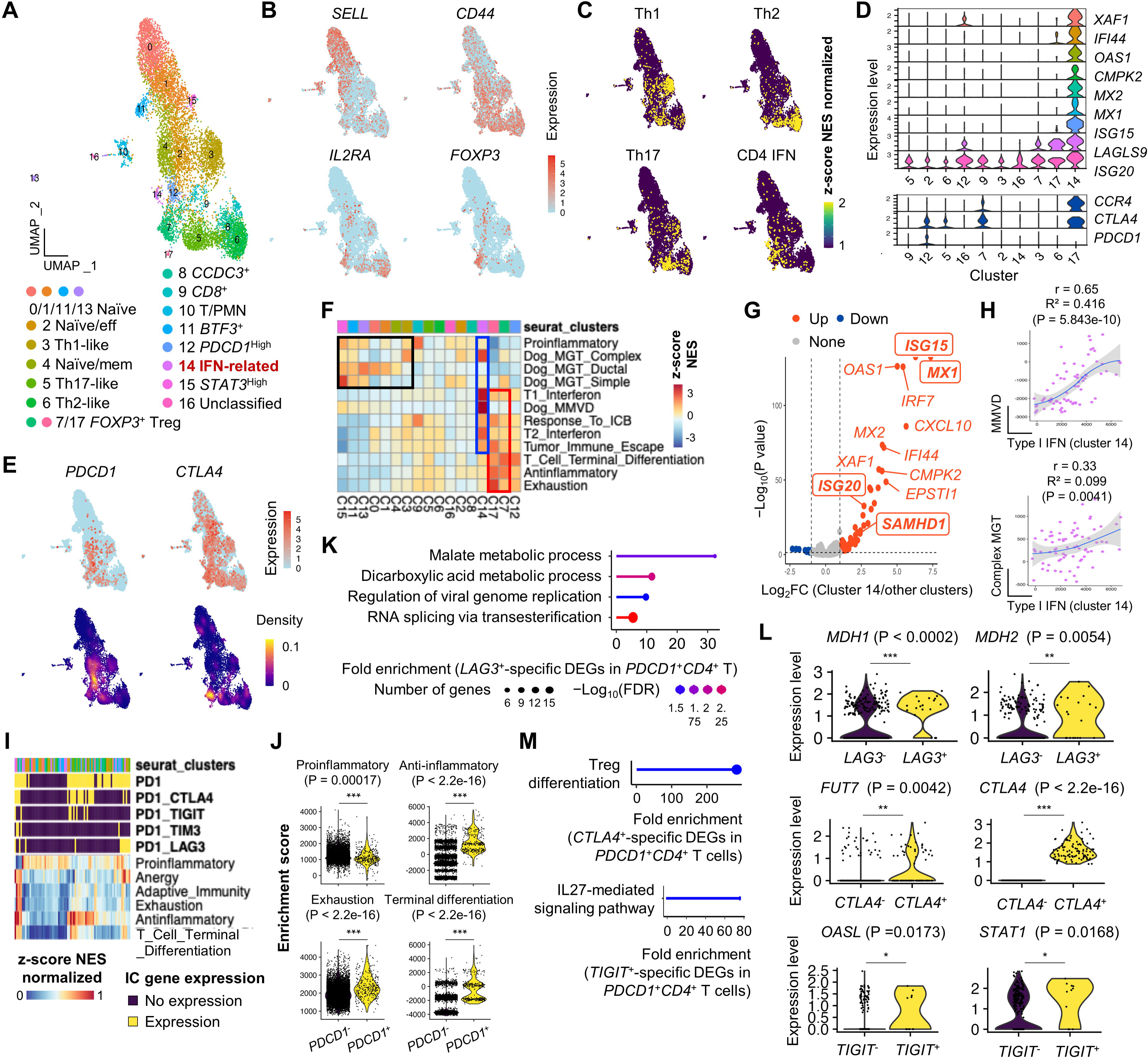
Identification and characterization of *CD4*^+^ T subpopulations by scRNA-seq. (A) UMAP plot showing 18 distinct *CD4*□ T cell subsets. (B) Cell type recognition using the escape R package. Feature plots show enrichment of canine Th1, Th2, and Th17 gene signatures along with *FOXP3* expression. (C) Expression patterns of immune checkpoint genes are shown on the feature plot. (D) Violin plots highlighting distinct gene expression in clusters 14 (IFN-related) and 12 (*PDCD1*□*CTLA4*□). (E) Feature and density plots showing the expression patterns of *PDCD1* and *CTLA4* in *CD4*□ T cells. (F) GSEA showing preferential pathway enrichment in effector *CD4*□ T phenotypes (red boxed). IFN-related cluster 14 is strongly enriched for IFN signaling and Dog_MMVD signatures. Treg and *PDCD1*^high^ clusters are enriched for terminal differentiation pathways. (G) Volcano plot showing DEGs in IFN-related *CD4*□ T cells compared to other clusters. Dog_MMVD-associated genes are highlighted in bold and boxed. (H) Feature scatter plots demonstrating positive correlation between two gene signatures in the IFN-related *CD4*^+^ T cells. (I) GSEA reveals that *CTLA4*□ or *LAG3*□*PDCD1*□ *CD4*□ T cells are enriched for anti-inflammatory, anergy, and exhaustion signatures. (J) Violin plots showing enrichment pattern of immune-associated gene sets in *CD4*□ T cells with or without *PDCD1*. (K) GO analysis of *LAG3*-associated DEGs in *PDCD1*□ *CD4*□ T cells reveals enrichment in malate and dicarboxylic acid metabolism, and viral genome replication. (L) Violin plots show upregulation of *LAG3*, *CTLA4*, and *TIGIT*-specific key genes involved in malate metabolism, Treg differentiation, and IL-27-mediated signaling pathways in *PDCD1*□*CD4*□ T cells, respectively. (M) GO analysis of *CTLA4*- and *TIGIT*-specific DEGs in *PDCD1*□*CD4*□ T cells reveals enrichment in Treg differentiation and IL-27-mediated signaling pathways, respectively. Statistical significance was determined by comparing two groups of interest using the non-parametric Wilcoxon rank-sum test. *P < 0.05, **P < 0.01, and **\***P < 0.001. Abbreviations: FC, fold change; DEG, differentially expressed genes; C, cluster; Treg, regulatory T cells; GSEA, gene set enrichment analysis; GO, gene ontology.

We next explored the functional and clinical relevance of these subsets. First, similar to myeloid cells, IFN-related *CD4*^+^ T cells exhibited marked enrichment of canine MMVD gene signature (**blue boxed in Figure 3F**), characterized by upregulation of *MX1* and *ISG15* (**Figure 3G**) and a positive correlation between type I IFN and MMVD signatures (**Figure 3H, upper panel**). A similar pattern was observed in the IFN-related subset regarding mammary complex carcinoma signature, showing upregulation of *ISG20* and *SAMHD1*, along with a corresponding gene set correlation (**Figures 3F, 3G, and 3H, lower panel**). Second, Treg clusters were highly enriched in anti-inflammatory signatures, consistent with their immunosuppressive roles in healthy dogs (49) (**red boxed in Figure 3F**). Interestingly, upon sub-clustering of Tregs based on *CCR4* expression (**Supplementary Figure 7C**), *CCR4*^+^ Tregs were more enriched with gene sets associated with tumor immune evasion and response to immune checkpoint (IC) blockade (**Supplementary Figure 7D**), supporting previous ex vivo findings in canine patients with genitourinary carcinomas (50,51). DEGs defining *CCR4*^+^ Tregs compared to *CCR4*^-^ ones were obtained and subject to GO analysis (**Supplementary Data 9**), which revealed significant enrichment with the gene set associated with tRNA wobble base modification (**Supplementary Data 10**). Among genes listed in the signature, *CCR4*^+^ Tregs showed a significant upregulation in *NSUN3*, *ELP2*, *GTPBP3*, *ELP3*, and *CTU1* but *ADAT2* downregulation. Third, *PDCD1*^high^*CD4*^+^ T cells showed modest enrichment of gene signatures associated with terminal T cell differentiation and exhaustion (**Figure 3F**), suggesting that healthy dogs have *CD4*^+^ T cells undergoing transition toward an exhausted phenotype. Finally, *CD4*^+^ T subsets enriched for mammary gland tumor (MGT)-associated gene signatures included several naïve, Th1- like, and *STAT3*^high^ proinflammatory populations (**black boxed in Figure 3F**).

In healthy dogs, PD-1 blockade has been shown to reverse CD4□ T cell suppression induced by tumor-derived PD-L1 (52), but the underlying molecular mechanisms are not well understood. We therefore analyzed the transcriptional profiles of *CD4*□ T cells expressing IC genes. *CD4*□ T cells with or without *PDCD1* expression exhibited different enrichment patterns for gene signatures associated with immune activation and exhaustion (**Figure 3I**). Notably, compared to *PDCD1*□*CD4*□ T cells, *PDCD1*□*CD4*□ T cells showed significant enrichment of exhaustion-related gene signatures, suggesting an inhibitory effect of *PDCD1* on *CD4*^+^ T cell immunity (**Figure 3J**). In the same context, *PDCD1*□*CD4*□ T cells showed downregulation of T cell receptor (TCR)-related genes (*ID3*, *CCR7*, *FOXP1*, and *TRAT1*) but also showed upregulation of genes such as *BATF*, *ISG20*, *IFI30*, *ITGB1*, and *IDH2* (**Supplementary Data 11**). Recently, a dog with anti-PD-L1-resistant melanoma exhibited a complete remission of an oral neoplastic lesion following anti-CTLA4 therapy (53). To further elucidate the mechanism of action, we performed differential gene expression and gene set enrichment analyses, which revealed that *CTLA4*-specific genes, such as *FOXP3*, *TIGIT*, *HAVCR2*, *IL10*, *ADORA2A*, and *ADA* (**Supplementary Figure 7E and Data 12**), were significantly upregulated and enriched in pathways related to negative regulation of the immune system (**Supplementary Figure 7F**). We next investigated the biological implications of additional IC genes expressed in *PDCD1*□*CD4*^+^ T cells. Among this population, distinct subgroups co-expressed additional IC genes, including *CTLA4* (n=95), *LAG3* (n=20), *TIGIT* (n=11), and *HAVCR2* (n=4). Notably, *PDCD1*□*CD4*□ T cells co-expressing *TIGIT*, *HAVCR2*, or *LAG3* showed a tendency toward enrichment of an exhaustion signature (**Supplementary Figure 7G**). DEGs associated with each IC gene are provided in **Supplementary Data 13**. Given the robustness of differential expression analysis in scRNA-seq datasets (54), GO enrichment analysis of *LAG3*-, *CTLA4*-, and *TIGIT*-specific DEGs was performed. Interestingly, *LAG3*-specific genes were significantly enriched in biological processes related to malate and dicarboxylic acid metabolism, viral genome replication, and RNA splicing (**Figure 3K**). Within the metabolic gene set, *MDH1* and *MDH2* were significantly upregulated in *LAG3*□*PDCD1*□*CD4*□ T cells compared to *LAG3*□ counterparts (**Figure 3L**). Similarly, *CTLA4*- and *TIGIT*-specific genes were enriched in pathways associated with Treg differentiation and IL-27 signaling, respectively (**Figure 3M**). In these gene sets, *FUT7*, *OASL*, and *STAT1* were notably upregulated in *IC*□*PDCD1*□*CD4*□ T cells (**Figure 3L**). Collectively, our results show the functional diversity and immune checkpoint heterogeneity of *CD4*□ T cell subsets in dogs, highlighting their potential clinical relevance in immune homeostasis.

### Identification and characterization of *CD8*^+^ and other T subsets

Next, we performed sub-clustering of non-*CD4*□ T cells (**Figure 4A**), revealing ten transcriptionally distinct *CD8*□ T cell clusters (0, 1, 2, 3, 4, 5, 6, 7, 8, and 14), three γδ T cell clusters (15, 17, 18), two cycling T clusters (9, 12), a double-negative (DN) *CD4*□*CD8A*□ T cluster (13), and unclassified T cell clusters (10, 11). Representative canonical and cluster markers used to annotate these T subsets are available in **Supplementary Figure 8A and Data 14**. Using unbiased cell type annotation based on established canine *CD8*□ T cell signatures (8,22,24), we identified several functional subsets (**Figure 4B**). Naïve *CD8*□ T cells (clusters 3 and 5) expressed *CCR7*, *SELL*, *LEF1*, and *TCF7*, while effector *CD8*□ T cells (clusters 0, 1, 2, 4, 6, 7, 8, and 14) showed high levels of cytotoxic genes including *GZMB*, *GZMH* (*LOC490629*), *GZMK*, and *PRF1* (**Figure 4C and Supplementary Figures 8A and 8B**). Clusters 2 and 6 expressed *FCER1G* and *PI3*, indicating an innate-like phenotype (9). Cluster 6 further expressed *CXCR6* and was enriched in effector/memory-associated markers, suggesting an effector memory mucosal-associated invariant T cell-like phenotype. Memory-like profiles were also observed in clusters 0, 8, and 14, based on the expression of *IL7R*, *ZNF683*, *PRDM1*, and *STAT3*. Cluster 7 comprised cells exhibiting high expression of *PDCD1* and *LAG3*. *CTLA4* expression was largely confined to cycling T (51.6% and 59.7% in clusters 9 and 12, respectively) and DN T cells (54.4% in cluster 13) in which cycling T subset was further characterized by *FOXP3*, *IL2RA*, *CTLA4*, and *IL10* expressions. γδ T cells were characterized by specific expression of *RHEX* and *SCART1* (*LOC491694*). Unclassified clusters (10 and 11) expressed *CD44*, *TOX*, and *LOC102155278*, while cluster 16, which expressed *CD8A*, *CSF3R*, and *DPYD*, likely represented doublets.

**Figure 4.**
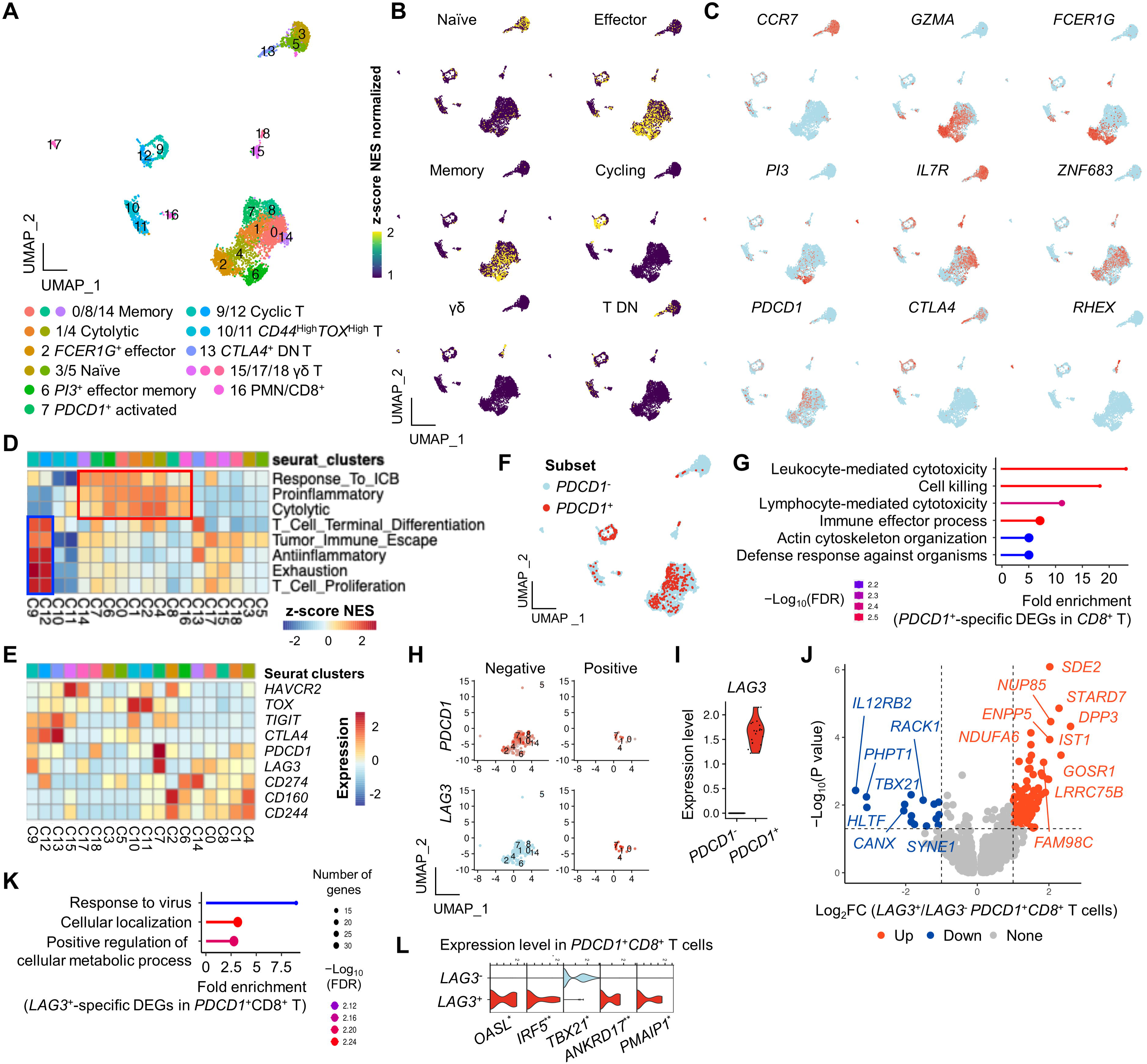
Identification and characterization of *CD8*^+^ and miscellaneous T subpopulations by scRNA-seq. (A) UMAP plot showing ten *CD8*□ T, three γδ T, two cycling T, and three unclassified T cell subsets. (B) Feature plots showing enrichment of canine gene signatures associated with naïve, effector, memory, cycling, γδ, and T DN T cells. (C) Feature plot of representative marker gene expression. (D) GSEA showing enrichment of immune-associated pathways across *CD8*□ T subsets. Effector phenotypes (red box) and cycling T subsets (blue box) are enriched in proliferation and exhaustion-related signatures. γδ T cells exhibit mild enrichment in naïve, exhausted, and proliferative signatures. Clusters 2, 4, and 7 show enrichment in cytolytic, proinflammatory, memory, and terminal differentiation pathways. (E) Heatmap of genes associated with T cell terminal differentiation. *PDCD1* and *LAG3* are highly expressed in cluster 7. (F) Feature plot showing *PDCD1*□ T cell distribution. (G) GO analysis of *PDCD1*-specific DEGs reveals enrichment of effector function and cytotoxicity-related biological processes. Feature (H) and violin (I) plots showing *LAG3*□*PDCD1*□*CD8*□ T cells. (J) Volcano plot of *LAG3*-specific DEGs in *PDCD1*□*CD8*□ T cells. (K) GO analysis of *LAG3*-specific DEGs in *PDCD1*□*CD8*□ T cells shows enrichment in antiviral response processes. (L) Violin plots of representative genes in the antiviral response pathway. All genes except *TBX21* are significantly upregulated in *PDCD1*□*CD8*□ T cells. Statistical significance was assessed using the Wilcoxon rank-sum test. *P < 0.05; **P < 0.01. Abbreviations: PMN, polymorphonuclear cells, FC, fold change; DEG, differentially expressed genes; C, cluster; T DN, double-negative T; GO, gene ontology.

Functional characterization using GSEA revealed several distinct patterns. First, effector *CD8*□ T cells were mainly enriched in proinflammatory, cytotoxic, and response to IC blockade gene sets (**red boxed in Figure 4D**), indicating functional competence, such as anti-tumor immunity in dogs (55). Second, T cell activation-associated gene signatures were markedly enriched in cycling T cells (**blue boxed in Figure 4D**), suggesting that cycling T cells may represent a state of steady-state antigenic stimulation (56). Third, although previous studies reported exhausted *CD8*□ T subsets in healthy dogs (9), our data showed that *PDCD1*□*CD8*□ T cells, despite expressing co-inhibitory receptors and being enriched for the T cell terminal differentiation signature (**Figure 4E**), did not exhibit remarkable enrichment of an exhaustion signature (**Supplementary Figure 8C**). Fourth, γδ T subsets also showed enrichment in T cell activation along with tumor immune escape signatures, suggesting a potential role in tumor immune surveillance. Lastly, although positioned adjacent to naïve T subsets on the dimensional plot, the DN T subset was enriched for anti-inflammatory and T cell terminal differentiation gene signatures, along with remarkable *CTLA4* and *TIGIT* expression. Additional pseudotime analysis revealed that the DN T subset was localized near the root of the trajectory and was not associated with any terminal lineage (**Supplementary Figures 8D**). Gene expression dynamics along eight terminal lineages revealed that *CTLA4* was transiently upregulated during early pseudotime in multiple lineages, particularly in cell fate 1 and 2 (**Supplementary Figures 8F**), suggesting DN T subset as an initial activation phase prior to differentiation.

To further explore the immunoregulatory role of *PDCD1* and co-inhibitory receptors in *CD8*□ T cells, we performed DEG analysis between *PDCD1*□*CD8*□ and *PDCD1*□*CD8*□ T cells (**Figure 4F**). *PDCD1*-specific DEGs (**Supplementary Data 15**) were enriched in immune effector pathways such as cytotoxicity, cell killing, and defense response (**Figure 4G and Supplementary Data 16**). Genes like *BATF, SH2D1A*, *CORO1A*, *PRF1*, and *GZMB* were commonly upregulated, while *IL7R* was downregulated. Meanwhile, similar to *CD4*^+^ T cells, *PDCD1* expression was significantly associated with an inhibitory effect on *CD8*^+^ T cell immunity (**Supplementary Figure 8F**). To further investigate functional implications of IC genes, *PDCD1*□*CD8*□ T cells were sub-grouped based on the expression of *LAG3* (n=20) (**Figures 4H and 4I**), *HAVCR2* (n=8), *CTLA4* (n=3), and *TIGIT* (n=4). Considering robustness of differential expression analysis in scRNA-seq datasets, we focused on *LAG3*-specific DEGs for downstream functional analysis (**Figure 4J and Supplementary Data 17**). GO enrichment revealed a significant association with viral response pathways (**Figure 4K**). Among the enriched genes, *PMAIP1*, *ANKRD17*, *IRF5*, and *OASL* were significantly upregulated, whereas *TBX21* was downregulated (**Figure 4L**). Meanwhile, *PDCD1*□*CD8*□ T cells with *CTLA4* and *LAG3* expression, showed a tendency for the enrichment of an exhaustion signature (**Supplementary Figure 8G**). Taken together, our results reveal transcriptionally distinct *CD8*□ T cell subsets in healthy dogs, with functional profiles supporting roles in immune homeostasis.

### Identification and characterization of B subsets

We performed sub-clustering of *CD79B*□ B cells and *MZB1*□ plasma cells, which identified ten distinct clusters (**Figure 5A**). Canonical and cluster-defining genes for each subset are shown in **Figures 5B and 5C**. Unbiased cell type annotation using canine B cell signatures classified the subsets into naïve B cells (clusters 0, 1, 2, 3, 6, and 9) and plasma cells (clusters 4, 5, and 8) (**Figure 5D**). Naïve B cells expressed *LOC100685971*, *CD79B*, *BTG1*, *MS4A1*, and MHC class II–associated genes, including *HLA*-*DRB1* and *DLA*-*DRA* (**Figure 5E**). Among them, clusters 2, 6, and 9 showed additional expression of *GNG2*, *VPREB3*, and *MX1*, and were enriched for type I IFN-related signatures (**Figures 5D and 5E**). Plasma subsets were marked by *JCHAIN*, *MZB1*, *PRDM1*, and *XBP1,* but lacked MHC class II expression, such as *DLA*-*DMB* and *DLA***-***DOA*. Notably, clusters 5 and 8 were transcriptionally defined by *UBE2C* and *OSBPL10*, respectively. The *UBE2C*□ plasma subset showed a high G2M cell cycle score, suggesting ongoing proliferation in steady-state (**Figure 5F**). Cluster 7 was excluded due to potential doublet identity based on *CD3E* and *DPYD* expression.

**Figure 5.**
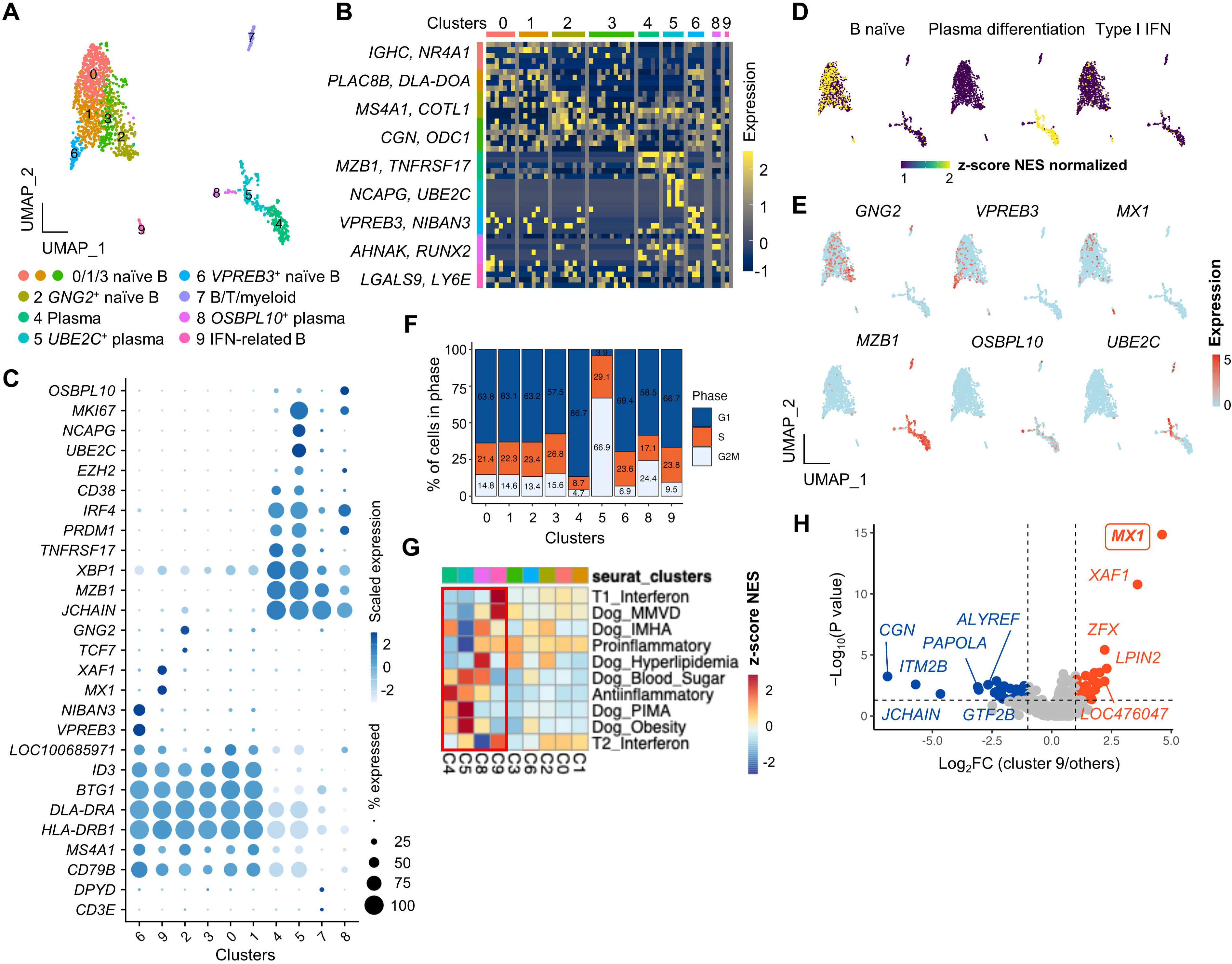
Identification and characterization of B subpopulations by scRNA-seq. (A) UMAP plot showing six B cell and three plasma cell subsets. (B) Heatmap presenting the top five DEGs defining each subset. (C) Feature plots showing enrichment of naïve and plasma cell gene signatures, along with representative marker expression. (D) Dot plot displaying canonical markers used to identify functionally distinct B cell subsets. (E) Cell cycle phase distribution across B cell subsets. (F) GSEA showing enrichment patterns of immune-related pathways across B cell subsets. (G) Volcano plot presenting DEGs in IFN-related B cells compared to other B cell subsets. MX1, listed in the Dog_MMVD signature, is highlighted in bold and boxed.

For functional characterization, GSEA revealed significant enrichment in plasma and IFN-related B subsets (**Figure 5G, red box**). Plasma subsets were enriched with gene sets associated with anti-inflammatory responses and canine immune/metabolic conditions, including IMHA, PIMA, obesity, blood sugar, and hyperlipidemia. The IFN-related B subset was highly enriched in canine MMVD signatures, with significant upregulation of *MX1* (**Figure 5H**). There was a weak positive correlation was observed between type I IFN and MMVD signatures (r = 0.309, P = 0.1723), but it did not reach statistical significance. Collectively, these findings reveal transcriptionally heterogeneous B and plasma cell subsets, with potential involvement in immune dysregulation and lipid metabolism.

### Cell-cell communication identifies key signaling pathways with ligand and receptor genes in regulating normal homeostasis in healthy dogs

Coordinated crosstalk among immune subsets is essential for maintaining immune homeostasis (31). To explore this, we investigated the cell-to-cell interactions involved in modulating immune equilibrium in healthy dogs. The overall signaling landscape across immune subsets is shown in **Supplementary Figure 9A**. In healthy dogs, we identified three major patterns of cell-cell interaction (**Figures 6A and 6B**).

**Figure 6.**
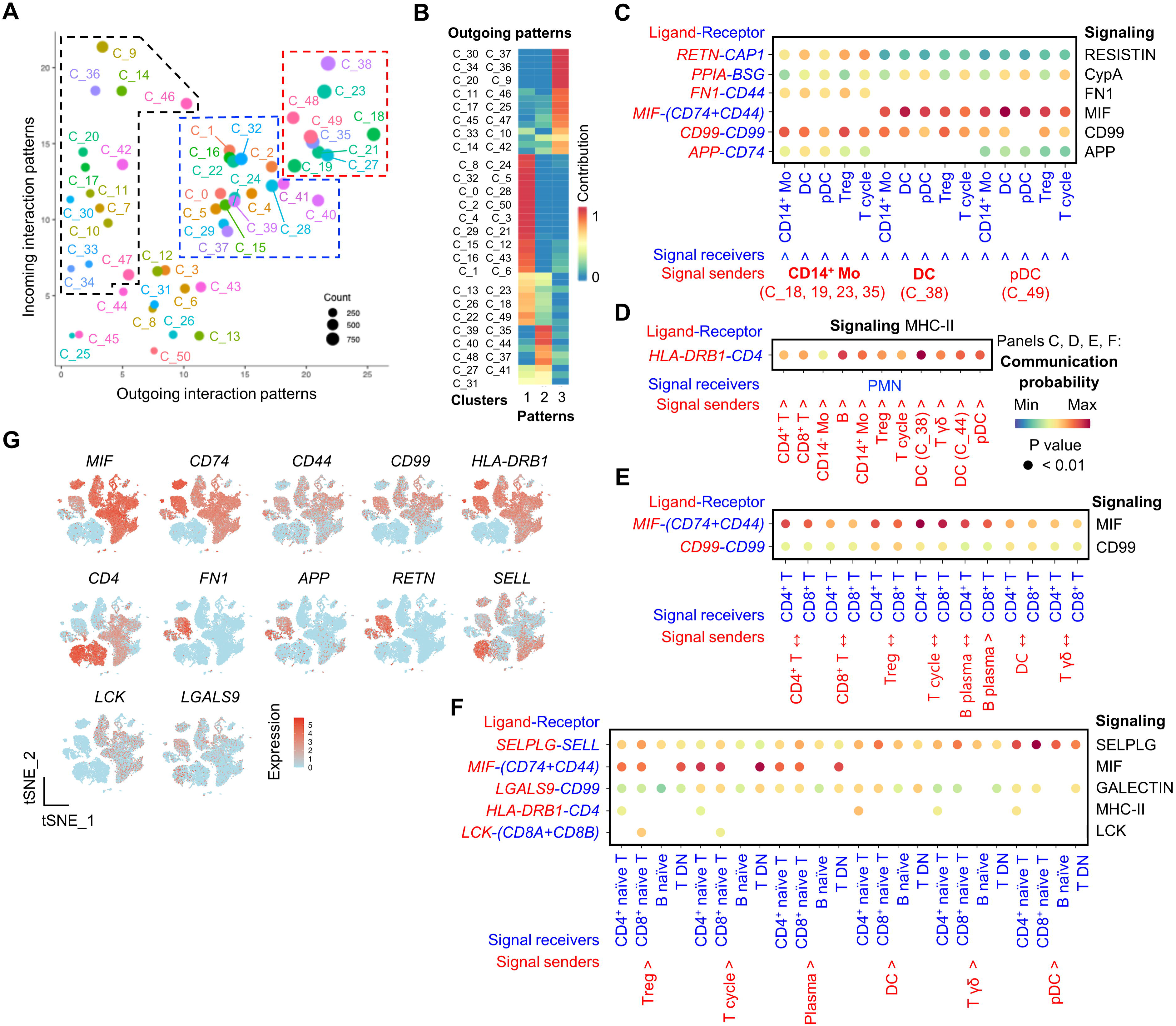
Identification of immune cell interactions by scRNA-seq using CellChat analysis. (A) Inferred intercellular communication network highlighting dominant signal senders (red and blue dashed lines) and receivers (black dashed line) visualized on a scatter plot. (B) Global outgoing signaling patterns across identified immune cell clusters. (C-F) Bubble plots illustrating significant ligand–receptor interactions involved in representative signaling pathways among indicated immune subsets. (G) Feature plots show the expression patterns of representative ligands and receptors involved in MIF, CD99, MHC-II, Selplg, FN1, APP, and Galectin signaling pathways.

First, autocrine signaling was strongly inferred among myeloid subsets, including DC (cluster 38), pDC (cluster 49), and *CD14*□ monocytes (clusters 18, 19, 23, and 35) (**Figure 6A, red dashed lines, Supplementary Figure 9A**). These myeloid subsets also transmitted signals to Treg (cluster 21), cycling T cells (cluster 27), and γδ T cells (cluster 48), suggesting a role in modulating T cell activity. The interactions involved secretory pathways (MIF, RESISTIN, CypA), cell-cell contact (CD99, APP), and extracellular matrix receptor interactions (FN1), through key ligand-receptor (LR) pairs such as *RETN*:*CAP1*, *FN1*:*CD44*, *MIF*:(*CD74*+*CD44*), and *CD99*:*CD99* (**Figure 6C**).

Second, neutrophils acted as primary signal recipients, prominently involved in the MHC-II signaling pathway (**Figure 6A, black dashed lines, Supplementary Figure 9B**). In this context, *CD4* was predicted to function as a key receptor, with *HLA*-*DRB1*□ subsets serving as ligand-bearing partners (**Figure 6D**). Third, effector immune subsets, including *CD4*□ T cells, *CD8*□ T cells, and plasma cells (clusters 0, 1, 2, 4, 5, 15, 16, 22, 24, 28, 29, 31, and 32), were predominantly engaged in MIF and to a lesser extent CD99 signaling. These involved the LR pairs MIF:(CD74+CD44) and CD99:CD99 (**Figures 6A, blue dashed lines, 6E, Supplementary Figures 9C and 9D**). Lastly, naïve and DN lymphocytes participated moderately in cell-cell contact (via SELPLG, LCK, and MHC-II) and secretory (GALECTIN, MIF) signaling pathways. Key LR genes included *SELL*, *CD44*, *CD74*, and *CD99* (**Figures 6A and 6F, Supplementary Figure 9A**). All identified LR genes are listed in **Figure 6G**. Taken together, these interactome analyses highlight key ligand-receptor axes and signaling pathways critical for immune crosstalk and homeostasis in healthy dogs.

### Integrated scRNA-seq analysis of circulating leukocytes in healthy dogs

Healthy dogs immunologically experience (57), exhibiting a broad and dynamic immune repertoire shaped by environmental exposures (9,58,59). To explore cohort-specific immune variation, we performed an integrated analysis of publicly available scRNA-seq datasets of peripheral leukocytes from two independent cohorts of healthy dogs (8,9). Using a standardized pipeline for preprocessing, quality control, data integration, and removal of potential doublets and dataset-origin bias (**Supplementary Figure 10A**), we analyzed 90,992 canine immune cells, ultimately identifying 35 transcriptionally distinct clusters in the integrated Seurat object (**Supplementary Figure 10B**). Cluster-defining marker genes are listed in **Supplementary Data 18.**

Integration revealed that immune cells from both cohorts generally retained a shared transcriptomic structure (**Supplementary Figure 10C**). However, proportional differences in immune subsets were observed between cohorts (**Supplementary Figure 10D**), such as clusters 16, 29, and 33 characterized by gene expression associated with platelets (*PPBP*), erythrocytes (*LOC100855558*, a hemoglobin subunit alpha-like gene), and B cells (*CD79B*), respectively (**Figure 7E**). One particularly study-specific subset, cluster 25, exhibited high expression of cell cycle-related genes (*PCLAF*, *MKI67*, and *SPC24*) and was enriched for a T cell cycling signature (**Supplementary Figures 10F and 10G**), suggesting a population of proliferating T cells potentially responding to steady-state antigenic stimuli (56). We further examined cohort-dependent expression of IC genes. Across studies, the proportions of *CTLA4*□ and *LAG3*□ T cells significantly differed (**Supplementary Figures 10H, 10I, Table 1**). Consistent with prior observations (60), *CTLA4*□ T cells comprised approximately 25% of circulating T cells in healthy dogs. Additionally, proportions of *PDCD1*□, *HAVCR2*□, *TIGIT*□ T cells, and *CD274*□ myeloid cells also differed by cohort. For example, *PDCD1*□ T cells were rarely observed in cohort by Ammons *et al*. (8), but represented 4.1% of T cells in cohort by Eschke *et al*. (9). These *PDCD1*□ T cells, associated with IC gene expression, were linked to immune activation, whereas *LAG3*□ and *TIM3*□*PDCD1*□ T cells were enriched for T cell exhaustion signatures (**Supplementary Figure 10J**). Importantly, T cell exhaustion enrichment scores differed significantly between studies (**Supplementary Figure 10K**), highlighting distinct immunological imprints shaped by environment, lifestyle, or microbial exposure. In summary, this integrated scRNA-seq analysis uncovers cohort-specific immune features in healthy dogs, providing evidence on how environmental context contributes to immune diversity and checkpoint-related immune phenotypes.

**Table 1.**
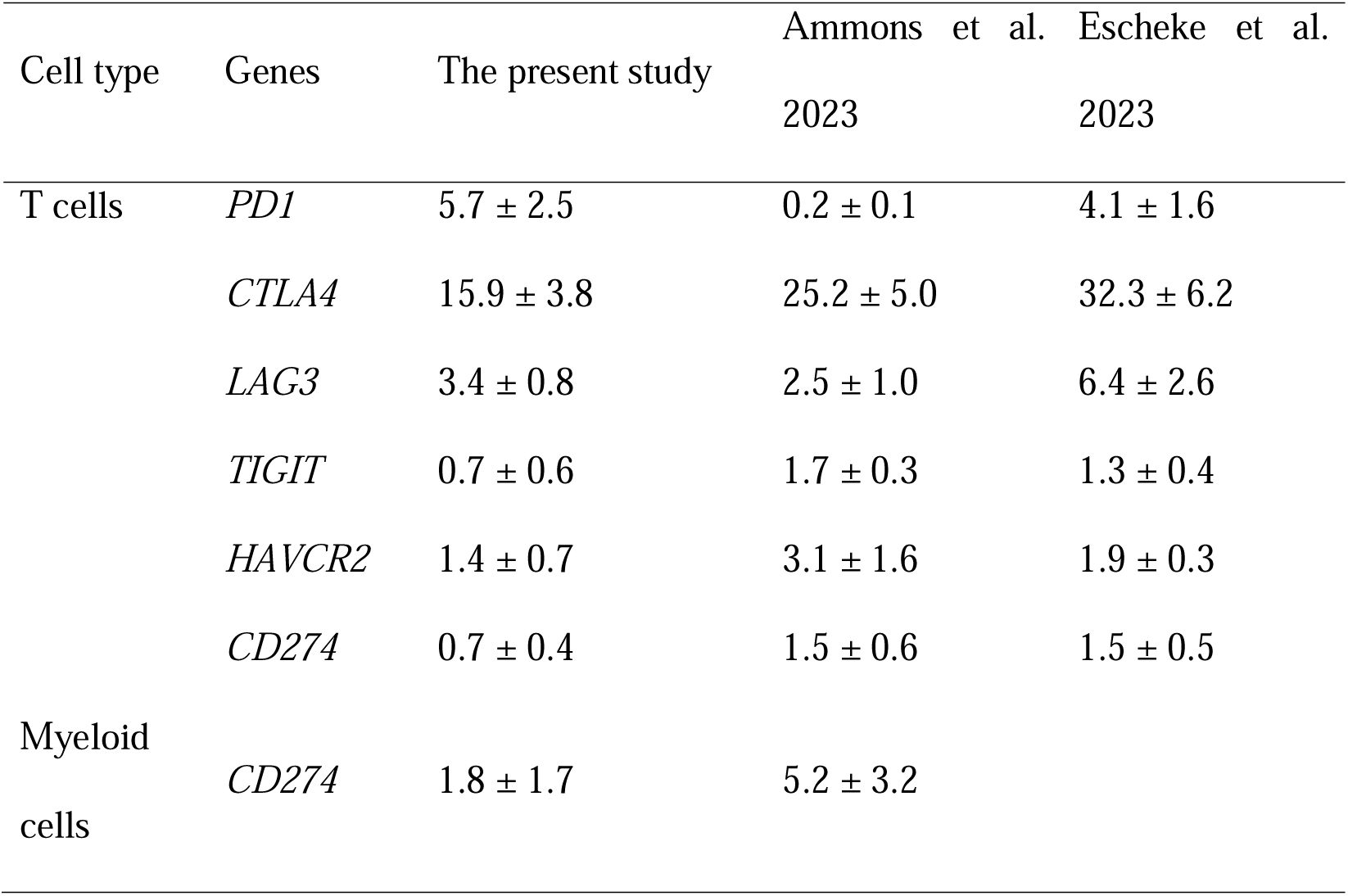
Proportions of immune subset with immune checkpoint genes across cohorts.

## Discussion

Companion dogs are recognized as valuable translational animal models for studying human diseases. Although emerging studies have applied scRNA-seq to canine peripheral leukocytes (8–10,13,16,20,23), the identification of functionally distinct and clinically relevant immune subsets, along with the elucidation of their roles and underlying molecular mechanisms in immune homeostasis, remains largely unexplored. To address the existing knowledge gap, we constructed a single-cell atlas of peripheral leukocytes from clinically healthy dogs adopting the canFam4 genome reference for scRNA-seq analysis for the first time. We demonstrate that the canFam4 provide a unique advantage over canFam3.1 by improving single-cell recovery and enabling the discovery of previously uncharacterized immune subsets with potential clinical relevance. To our knowledge, this is also the first study to provide mechanistic insights into how immune checkpoint genes, such as *PDCD1*, *CD274*, and *LAG3*, contribute to normal immune homeostasis in healthy dogs. Finally, we demonstrate that environmental context has great potential to shape cohort-dependent immune phenotypes and diversity. Collectively, our dataset and analytical framework offer a valuable resource for translational immunology in dogs, with implications for cancer research and beyond.

canFam3.1 has traditionally served as the main reference for sequence annotation in dogs, but recent studies have begun to adopt canFam4 for transcriptomic profiling of canine immune cells (61,62) due to its superior contiguity and enhanced transcriptional resolution (30). In the present study, we applied canFam4 to annotate single-cell transcriptomic data derived from circulating leukocytes in healthy dogs. For the first time, our results demonstrated that canFam4 could not only improve transcriptomic resolution by identifying previously unannotated gene features but also, of note, increase the number of assignable cells while reducing the proportion of reads mapped to non-human homologous genes. These improvements are believed to facilitate the discovery of novel lineage-specific and functionally relevant features, thereby enabling more precise identification and characterization of transcriptionally heterogeneous immune cell subsets in dogs. For example, *CD4*□ monocytes have been suggested to resemble *CD14*□ monocytes in dogs (8). However, our data revealed that *CD14*□ monocytes rarely expressed *CD4*, which was instead mainly detected in neutrophils, consistent with prior flow cytometric findings (63). Beyond known markers such as *CD1c*, *CD86*, *CD83*, and *IL3RA*, we identified additional genes, such as *CD11c* (*ITGAX)*, CD1 family of genes (*CD1A*, *CD1B*, *CD1D*, and *CD1E*), *HLA*-*DRB1*, *DLA*-*DQB1*, *DLA*-*64,* and *CD33*, which could serve as canonical markers for DC. Similarly, *CD21*, *CD24*, and *CEACAM1* could prove useful for identifying low-density granulocytes or MDSCs. For T cells, *GLNY*, *NKG7*, *GZMH*, *IL32*, and *BATF* may assist in delineating effector *CD8*□ T and *CD4*□ Treg subsets in dogs. Currently, canine genome references remain incompletely annotated (10). Thus, the use or integration of updated genome assemblies, such as canFam4, may not only offer great potential to uncover the clinical relevance of previously uncharacterized immune subsets (8,10,64), but also provide a strategic approach to enhance the analytical power of single-cell transcriptomics in canine immunological research. For example, *IL32* and *BATF* have been implicated in the immunosuppressive functions of tumor-infiltrating Tregs (TI-Tregs) (65,66). Thus, investigating the *IL32*□*BATF*□ Treg subset may shed light on the molecular mechanisms by which canine TI-Tregs exert potent immunosuppressive effects and contribute to a tumor-permissive microenvironment. Likewise, *CEACAM1* and *CD24* have been shown to mediate neutrophil-driven immunosuppression during microbial infections and in cancer (67,68). CEACAM1 has been also implicated in T cell tolerance and exhaustion via interaction with the checkpoint receptor TIM3 (69). Interestingly, dogs have circulating CEACAM1□ granulocytes (70). In this context, investigating the role of cell-cell interactions mediated by canine CEACAM1□CD24□CD4□ granulocytes may be of particular interest, given their potential involvement in both microbial susceptibility and immune suppression within the tumor microenvironment. We also, for the first time, report that dogs have peripheral *XCR1*□ DC subset, consistent with findings in humans, mice, and cattle (71–73). As cDC1, XCR1□ DCs play a key role in rejuvenating exhausted T cell subsets during chronic viral infection and tumor progression (71,74). Future studies would be valuable to validate their existence in inflamed tissues and to assess their functional competence, which may inform the development of DC-targeted vaccines in dogs.

In this study, we identified several immune subsets that may help elucidate immunological mechanisms associated with disease pathogenesis. For example, IFN-enriched *CD14*□ monocytes and *CD4*□ effector T subsets showed significant enrichment of both type I IFN and MMVD-related gene signatures, along with upregulated expression of canonical IFN-stimulated genes, such as *ISG15* and *MX1*. Similarly, IFN-related B cell subset, which was first identified in our scRNA-seq analysis, exhibited enriching tendency to MMVD while significantly elevating *MX1* expression. Our results strongly indicate the potential involvement of IFN-related subset in the pathogenesis and disease mechanism of MMVD in dogs. Indeed, type I IFN signaling has been closely implicated in the progression of canine MMVD, with significant *MX1* upregulation in the valvular tissues of severely affected dogs (75). Reduced expression of IFN-related genes, including *ISG15* and *MX1*, has also been proposed to increase susceptibility to viral infections in the early stages of MMVD (44). In humans, these genes play a protective role in cardiac tissue against viral invasion (44). Of note, immune subsets with constitutive expression of IFN-stimulated genes, such as *MX1*, are considered pre-activated population that can mount rapid responses to malignant cells, pathogens, or autoantigens (31,76–78). In dogs, IFN-related *CD4*□ T and *CD14*□ monocyte subsets are thought to patrol and infiltrate inflamed tissues (8,22,24). Thus, based on our findings, we propose that IFN-enriched immune subsets are activated by MMVD-associated autoantigens, initiating aberrant immune responses that contribute to disease progression. The enrichment of *CD177*□ neutrophils in both MMVD and IMHA signatures further supports our hypothesis, suggesting a potential link between innate immunity and disease pathogenesis. *CD177*, a marker of activated neutrophils (32), is significantly upregulated in peripheral leukocytes from dogs with MMVD, along with *MMP9* and *CAMP*, which are key molecules implicated in myocardial remodeling in dogs (79) and cardioprotective responses in humans (80). Similarly, dogs with IMHA show significant upregulation of *PTX3*, *CP*, *C3*, *CAMP*, *MMP8*, *MMP9*, and *PGLYRP1* (35), many of which are representative DEGs defining the *CD177*^+^ neutrophil subset identified in our study. Given that mitral valves from dogs with MMVD could lack overt immune cell infiltration (81), we postulate that circulating soluble and/or paracrine mediators derived from *CD177*^+^ neutrophils contribute to the development of MMVD. Indeed, *CD177* (82), *PTX3* (83), and *C3* (84) have been implicated in promoting the release of neutrophil extracellular traps, which have been associated with the pathogenesis of canine IMHA (85,86). Collectively, our results provide a rationale for future longitudinal single-cell and secretome profiling of blood samples from dogs with either progressive MMVD or those undergoing surgical intervention. Such efforts may help elucidate MMVD mechanisms driven by dysregulated innate immunity and soluble factor-mediated pathways from IFN-related and *CD177*^+^ subsets.

As in humans, naturally occurring cancers are highly immunogenic in dogs (87,88). Checkpoint molecules play a pivotal role in shaping anti-tumor responses, determining the clinical outcomes of canine cancer patients (46,50,89,90). To date, however, biological functions and molecular mechanisms of the checkpoint genes, such as *PDCD1*, *CD274*, *LAG3*, and *CTLA4*, remain largely unexplored, which was investigated in this study by leveraging scRNA-seq. First, consistent with previous findings (52), our results demonstrate that *PDCD1* has inhibitory effects on T cell immunity in dogs. It should be noted that although genes and signaling pathways associated with T cell activation were upregulated in *PDCD1*^+^*CD4*^+^ and *CD8*^+^ T cells, those related to TCR signaling, such as *ID3*, *CCR7*, *FOXP1*, *TRAT1*, and *IL7R* were downregulated. This suggests that *PDCD1* should be interpreted as both an activation- and exhaustion-associated marker. Second, consistent with human data, *LAG3* was implicated in canine T cell exhaustion. For example, *LAG3*^+^*PDCD1*^+^*CD8*^+^ T cells exhibited downregulation of *TBX21*, which encodes T-bet, a transcription factor known to repress LAG3 expression to sustain a virus- or tumor-specific PD1^+^CD8^+^ T cell responses (91,92). In addition, *LAG3* was associated with upregulation of *MDH1* and *MDH2*, implicating a role for malate metabolism in *PDCD1*^+^*CD4*^+^ T cells. Malate metabolism mediated by MDH1 and MDH2 is essential for preserving the function of exhausted LAG3^+^PD1^+^CD8^+^ T cells during lymphocytic choriomeningitis virus infection (93). Future studies are warranted to determine whether malate metabolism directly modulates the functional stability of CD4^+^ T cells and prevents their transition toward exhaustion, which remains to be demonstrated in both humans and dogs. Third, we found that *CD274*^+^ neutrophils and monocytes may be involved in IL-10 production and PD-L1-mediated immunosuppression in cancer. In dogs, IL-10 has been shown to inhibit neutrophil function during babesiosis (94). In humans, tumor-infiltrating PD-L1^+^ neutrophils represent a major source of IL-10 production in lung cancer patients (95), and PD-L1 expression on CD14^+^ monocytes infiltrating hepatocellular carcinomas also contributes to IL-10 production (96). These findings raise the possibility that PD-L1^+^ neutrophils or PMN-MDSC in dogs may contribute to an immunosuppressive tumor microenvironment through IL-10 production, which warrants further investigation. Finally, consistent with previous findings (53,60,97), we found that *CTLA4* may contribute to immune tolerance in canine T subsets, including well-characterized Tregs, and notably, the less-defined DN T cells. In our study, the DN T subset expressed *CTLA4*, but lacked other previously reported regulatory markers, such *FOXP3*, *IL10*, *IL2RA*, and *GATA3* (98,99). Furthermore, given that this subset appears to originate from naïve T cells, and that neither trajectory nor GSEA analyses supported an exhausted phenotype, we propose that *CTLA4* may function as an early regulatory marker guiding the differentiation of DN T cells in dogs. Although direct evidence of DN T subset within canine tumors is currently lacking, their immunoregulatory properties in canine tissues (99,100) and the observed anti-tumor efficacy of CTLA4-targeted therapies in dogs (53) suggest that DN T cells may also play a functional role within the tumor microenvironment.

PBMCs reflect peripheral immunity, which can be influenced by various factors, such as breed, lifestyle, and environmental conditions (101–103). In healthy dogs, circulating T and myeloid cells express PD1, PD-L1, and CTLA4 (53,104,105). Herein, we investigated the cohort-specific effects on PBMC immune profiles by integrating publicly available scRNA-seq datasets from healthy dogs. For the first time, our integrated scRNA-seq analysis delineated cohort-specific immune profiling, as evidenced by differences in the proportions of cycling T cells, checkpoint-expressing immune subsets, and T cell exhaustion states. In dogs, cycling T cells may contribute to immune homeostasis in the periphery, duodenum, Peyer’s patches, and mesenteric lymph nodes, reflecting responses to antigenic stimulation (8,14,15,22). This notion is further supported by our interactome analysis, which reveals that cycling T cells interact with *CD14*^+^ monocytes, DC, or pDC via MIF and CD99 signaling pathways. Furthermore, each cohort displayed a distinct pattern of IC gene expression, such as *CTLA4* and *LAG3*, along with varying degrees of T cell exhaustion, underscoring the immune heterogeneity among individuals. These differences likely reflect cohort-specific antigenic stimulation. PBMC datasets have been widely shared as critical reference data not only for investigating naturally occurring diseases (8,106,107) but also for monitoring the impacts of environmental and lifestyle factors on immunity (108–111). Accordingly, it may be preferable to adopt immune cell reference datasets that are tailored to the local context, such as domestic conditions in South Korea, and the specific research purpose.

Finally, we investigated normal homeostatic cell–cell interactions among peripheral immune subsets in healthy dogs. Our interactome analysis demonstrates that MIF, CD99, MHC-II, and SELPLG represent key intrinsic cellular interplay in regulating immune homeostasis. Interestingly, Moore *et al*. (112) previously proposed that CD4 on canine neutrophils engage in molecular interactions with leukocyte-expressed receptors, which, in our study, were predicted to include *HLA*-*DRB1*. Notably, human and canine neutrophils endogenously express surface CD4, which has been shown to bind certain viral proteins of clinical relevance in promoting inflammation (113). Furthermore, CD4 expression on neutrophils has been associated with their migratory capacity toward sites of inflammation (114). In line with these findings, our study observed that *CD4*^+^ neutrophils were strongly enriched in inflammatory transmigration pathways, suggesting that MHC-II signaling may regulate the biodistribution of canine *CD4*^+^ neutrophils through interaction with *HLA*-*DRB1*^+^ leukocytes. Additionally, canine macrophages infiltrating local and metastatic oral melanoma were previously reported to significantly upregulate *CD99* and *MIF* compared to normal mucosal tissues (115). Blood levels of these molecules have also been found elevated in dogs with atopic dermatitis and mammary neoplasia (116,117). In our analysis, *CD14*^+^ monocytes and DCs showed homotypic interactions mediated by CD99 and MIF signaling pathways. Canine *XCR1*^+^ DC showed specific MIF expression. Such homotypic myeloid interactions have been implicated in shaping the anti-tumor immune response in human melanoma patients (118). Thus, we postulate that CD99 and MIF signaling may serve as critical components of homeostatic immune interactions, not only in the steady state but also under inflammatory and tumor microenvironmental conditions. These warrant further validation using single-cell and spatial transcriptomic approaches.

In conclusion, this is the first study to map the single-cell transcriptomic landscape of circulating leukocytes in South Korean dogs. Our results provide a valuable resource to support future scRNA-seq studies on immune cells in dogs affected by various immunological disorders. Methodology used in this study may help pave the way for investigating potential roles of immune checkpoint genes in translational research, bridging canine and human immunology.

## Acknowledgments

We thank the University of Florida High-Performance Computing Center for providing access to the HiPerGator 3.0 supercomputer, which served as the primary computational platform for integrating multiple scRNA-seq datasets in this study. We are also grateful to Dr. Weizhou Zhang for sponsoring Dr. Myung-Chul Kim’s use of HiPerGator.

## Data availability

The raw and processed scRNA-seq data generated in this study have been deposited in the NCBI GEO under accession number GSE301630. The raw data from scRNA-seq in this study was publicly available as described in the main manuscript with the accession numbers.

## Additional information Funding

This research was supported by the Basic Science Research Program through the National Research Foundation of Korea (NRF) funded by the Ministry of Education (No. 202300241779). This research was supported by the Bio & Medical Technology Development Program of the NRF funded by the Korean government (No. 2022M3A9D3016848).

## Author’s contribution

Conception and design: Myung-Chul Kim; Development of methodology: Myung-Chul Kim; Data acquisition: Myung-Chul Kim, Taeeun Gu, Hyeewon Seo, Yewon Moon, Hyun Je Kim; Bioinformatics: Myung-Chul Kim, Nicholas Borcherding; Analysis and interpretation of data: Myung-Chul Kim, Nicholas Borcherding, Ryan Kolb, Weizhou Zhang; Writing, review, and/or revision of the manuscript: Myung-Chul Kim, Nicholas Borcherding, Ryan Kolb, Weizhou Zhang, Youngmin Yun, Woo-Jin Song, Chung-Young Lee, Hyun Je Kim; Study supervision: Myung-Chul Kim

## Supplementary Figure Legends

**Supplementary Figure 1.** Preprocessing and quality control of the scRNA-seq data in Seurat. (A) Violin plots showing the distribution of unique genes detected per cell (nFeature_RNA), the percentage of mitochondrial reads, and the percentage of non-human homologous genes across biological replicates. (B) Selection of 3,000 highly variable features exhibiting high cell-to-cell variation within the integrated dataset. (C) PCs enriched for informative features (low p-values) are visualized using a JackStraw plot. (D) tSNE plot displaying immune cells from each biological replicate, annotated with cell counts. (E) tSNE distribution of immune cells by breed, age, and sex. No notable differences were observed across these groups. (F) Cell proportions across clusters and biological replicates. (G) Distribution and proportion of potential doublets detected in the integrated Seurat object. (H) Sub-clustering of predicted doublets, revealing co-expression of T, B, and myeloid lineage markers on the tSNE feature plot. Abbreviations: CM, castrated male; IM, intact male; SF, spayed female; PC, principal component.

**Supplementary Figure 2.** Canonical marker expression profiles used to define distinct single-cell immune clusters. Dot plot showing representative canonical markers used to annotate transcriptionally and functionally distinct immune subsets within the integrated scRNA-seq dataset.

**Supplementary Figure 3.** Unbiased cell type recognition using reference-based annotation tools. (A) Heatmap showing cell type predictions using the escape R package, based on canine immune cell gene signatures derived from previous studies (8,22). (B) tSNE plot showing cell type annotations performed using the SingleR R package with the Monaco Immune Cell Data as the reference dataset.

**Supplementary Figure 4.** Cell cycle analysis of immune cell subsets. Cell cycle phase distribution (G1, S, G2/M) is shown across immune cell subsets within the integrated scRNA-seq dataset.

**Supplementary Figure 5.** Preprocessing and comparative analysis of scRNA-seq data using canFam3.1 and canFam4 reference genomes. (A) Quality control metrics across the two canine genome references, including the number of cells per sample, number of unique genes detected per cell, percentage of non-human homologous genes, and percentage of mitochondrial reads. Each dot represents a biological replicate. (B) tSNE plot displaying 48 clusters identified from the integrated dataset aligned to canFam3.1. (C) tSNE density plots comparing the distribution of CD8□ T cell subsets between canFam3.1 and canFam4 references. (D) Heatmap showing representative genes among the top 10 DEGs used to define each major immune lineage. Cell counts per lineage are indicated below the heatmap. A full list of highly variable features specific to canFam4 compared to canFam3.1 is provided in **Supplementary Data 4**.

**Supplementary Figure 6.** Identification and characterization of myeloid subpopulations by scRNA-seq. (A) Dot plot showing canonical marker expression profiles used to define transcriptionally and functionally distinct myeloid cell subsets. Cluster annotations correspond to those in Figure 2. Violin plots displaying the expression pattern of *MIF* (B) and *LGALS9* (C) on myeloid subsets. (D) Feature plots displaying the expression patterns of *CD4*, *ITGAX*, and *CEACAM1* in myeloid cells. (E) Scatter plot showing the correlation between MMVD and IMHA signatures in cluster 20. (F) Volcano plot showing representative DEGs in IFN-related monocytes compared to other monocyte subsets. (G) Violin plot illustrating the segregation of myeloid subsets based on *CD274* expression. (H) Volcano plot displaying representative DEGs in *CD274*□ monocytes relative to *CD274*□ monocytes. *CD274* is highlighted with a log□FC of 12.8 and P = 0. (I) GO analysis of *CD274*□ monocyte-specific DEGs reveals enrichment in biological processes related to the PD-L1 checkpoint pathway in cancer. Abbreviations: FC, fold change; DEG, differentially expressed genes; GO, gene ontology.

**Supplementary Figure 7.** Identification and characterization of *CD4*^+^ T subpopulations by scRNA-seq. (A) Dot plot showing canonical markers used to identify functionally distinct CD4□ T cell subsets. (B) Feature plots showing expression patterns of *BATF* and *IL32*. (C) Feature plot showing expression patterns of *FOXP3*, *IL2RA*, *CTLA4*, and *CCR4* in Tregs sub-clustered from clusters 7 and 17. (D) Violin plots illustrating increased enrichment of gene sets associated with tumor immune escape and response to immune checkpoint blockade in *CCR4*□ Tregs. (E) Volcano plot showing representative *CTLA4*-specific DEGs that belong to the gene set of negative regulation of immune system. (F) GO analysis of *CTLA4*□*CD4*^+^ T-specific DEGs. (G) Violin plots showing increased enrichment of exhaustion gene set in *HAVCR2*□ and *CTLA4*□ *PDCD1*□*CD4*□ T cells. Statistical significance and P values was determined by comparing two groups of interest using the non-parametric Wilcoxon rank-sum test in the ggpubr R package.

**Supplementary Figure 8.** Identification and characterization of *CD8*^+^ T and miscellaneous subpopulations by scRNA-seq. (A) Canonical marker expression profile used to define distinct CD8□ and miscellaneous T cell clusters. Representative canonical markers are shown in the dot plot. (B) Feature plots showing the expression of *CD247* (*CD3Z*), *CD4*, and *CD8A*. (C) GSEA presenting immune-associated pathway enrichment in *CD8*□ T cell clusters (clusters 0, 1, 2, 3, 4, 5, 6, 7, 8, and 14) with immune checkpoint expression. (D) Slingshot-based pseudotime trajectory of *CD8*□ cells projected onto UMAP space. Red and blue dots refers to terminal and root nodes, respectively. Cells with gray indicate unassigned pseudotime. (E) *CTLA4* expression dynamics along eight terminal lineages or cell fates. Black lines represent smoothed local regression, and red dots indicate individual cell measurements. (F) Violin plots showing enrichment pattern of immune-associated gene sets in *CD8*□ T cells with or without *PDCD1*. (G) Violin plots showing increased enrichment of exhaustion gene set in *CTLA4*□ and *LAG3*□ *PDCD1*□*CD8*□ T cells.

**Supplementary Figure 9.** Inference of immune cell communication network by CellChat analysis. (A) Overall outgoing and incoming signaling patterns of significant pathways across clusters. Bar plots represent the total interaction strength for each sending (column) and receiving (row) cell type. (B-D) Network centrality analysis identifies the dominant signal senders, receivers, mediators, and influencers in each indicated signaling pathway.

**Supplementary Figure 10.** Integrated scRNA-seq analysis reveals cohort-specific immune features in healthy dogs. (A) Study scheme showing the overview of 20 healthy canine PBMC scRNA-seq datasets used in this study. (B) UMAP plot of the integrated Seurat dataset comprising 90,992 single cells classified into 35 clusters. Canonical immune markers used for major subset identification are indicated. (C) UMAP plot displaying cell distribution across different canine cohorts. (D) Bar plot showing the relative abundance of immune subsets across datasets, (E) Violin plots of representative genes enriched in specific subsets identified in the GSE225599 dataset. (F) Proportional analysis highlighting a study-specific cycling T subset (cluster 25). (G) Feature plot indicating T cell cycle gene signatures and proliferation-related genes in cycling T cells. (H) Comparative analysis of immune checkpoint gene–expressing T cell proportions, showing cohort-specific presence of *CTLA4*□ and *LAG3*□ cells. (I) Dimensional plot displaying the spatial distribution of T cells expressing immune checkpoint genes (*CTLA4*, *LAG3*), visualized in the front layer. (J) GSEA results indicating enriched immune-related pathways in T cells expressing immune checkpoint genes. P values were obtained using one-way ANOVA via the ggpubr R package and Wilcoxon rank-sum tests. * *P* < 0.05, ** *P* < 0.01, and *** *P* < 0.001.

## Supplementary Data Legends

**Supplementary Data 1.** Demographic characteristics of enrolled dogs.

**Supplementary Data 2.** Complete blood count and serum biochemical test results.

**Supplementary Data 3.** Summary of scRNA-seq library construction and multiplexing statistics.

**Supplementary Data 4.** Highly variable gene features identified for canFam3.1 and canFam4 genomes.

**Supplementary Data 5.** Top 20 cluster-defining genes for each myeloid cell subset.

**Supplementary Data 6.** *CD274*-specific DEGs in myeloid cells.

**Supplementary Data 7.** GO terms enriched from genes listed in Supplementary Data 6.

**Supplementary Data 8.** Top 20 cluster-defining genes for each *CD4*□ T cell subset.

**Supplementary Data 9.** *CCR4*-specific genes identified in Tregs.

**Supplementary Data 10.** GO terms enriched from genes listed in Supplementary Data 9.

**Supplementary Data 11.** *PDCD1*-specific genes in *CD4*□ T cells.

**Supplementary Data 12.** *CTLA4*-specific genes in *CD4*□ T cells.

**Supplementary Data 13.** DEGs associated with co-inhibitory receptors in *PDCD1*□*CD4*□ T cells.

**Supplementary Data 14.** Top 10 cluster-defining genes for each *CD8*□ and miscellaneous T cell subset.

**Supplementary Data 15.** *PDCD1*-specific genes in *CD8*^+^ T cells.

**Supplementary Data 16.** GO terms enriched from genes listed in Supplementary Data 15.

**Supplementary Data 17.** DEGs associated with co-inhibitory receptors in *PDCD1*□*CD8*□ T cells.

**Supplementary Data 18.** Top 20 cluster-defining genes in the integrated scRNA-seq dataset.

