## Supplementary figures and images for "High-resolution single-cell RNA sequencing using canFam4 reveals novel immune subsets and checkpoint programs in healthy dogs"

### Suppl_Figure_1.TIF

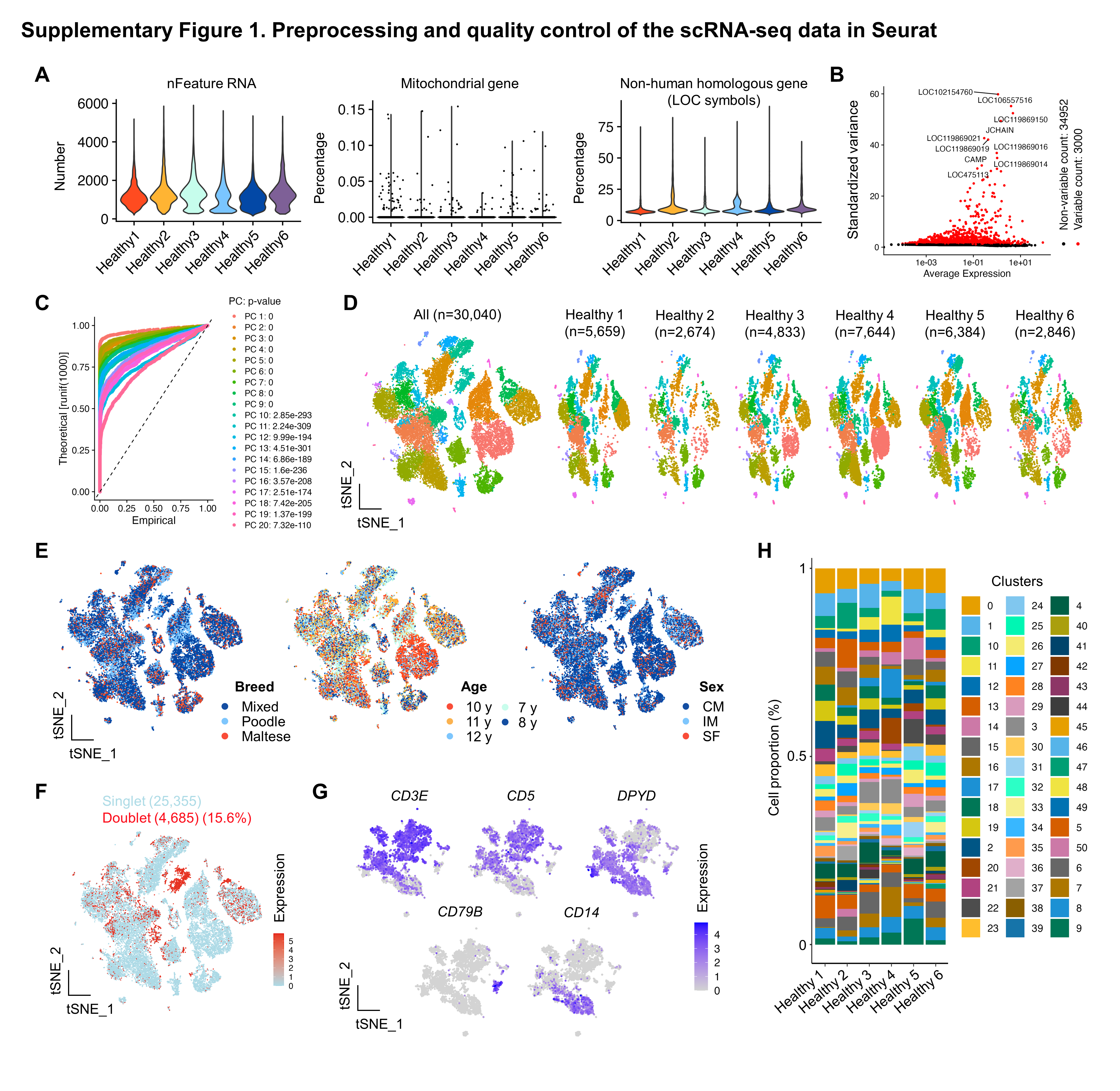

### Suppl_Figure_2.TIF

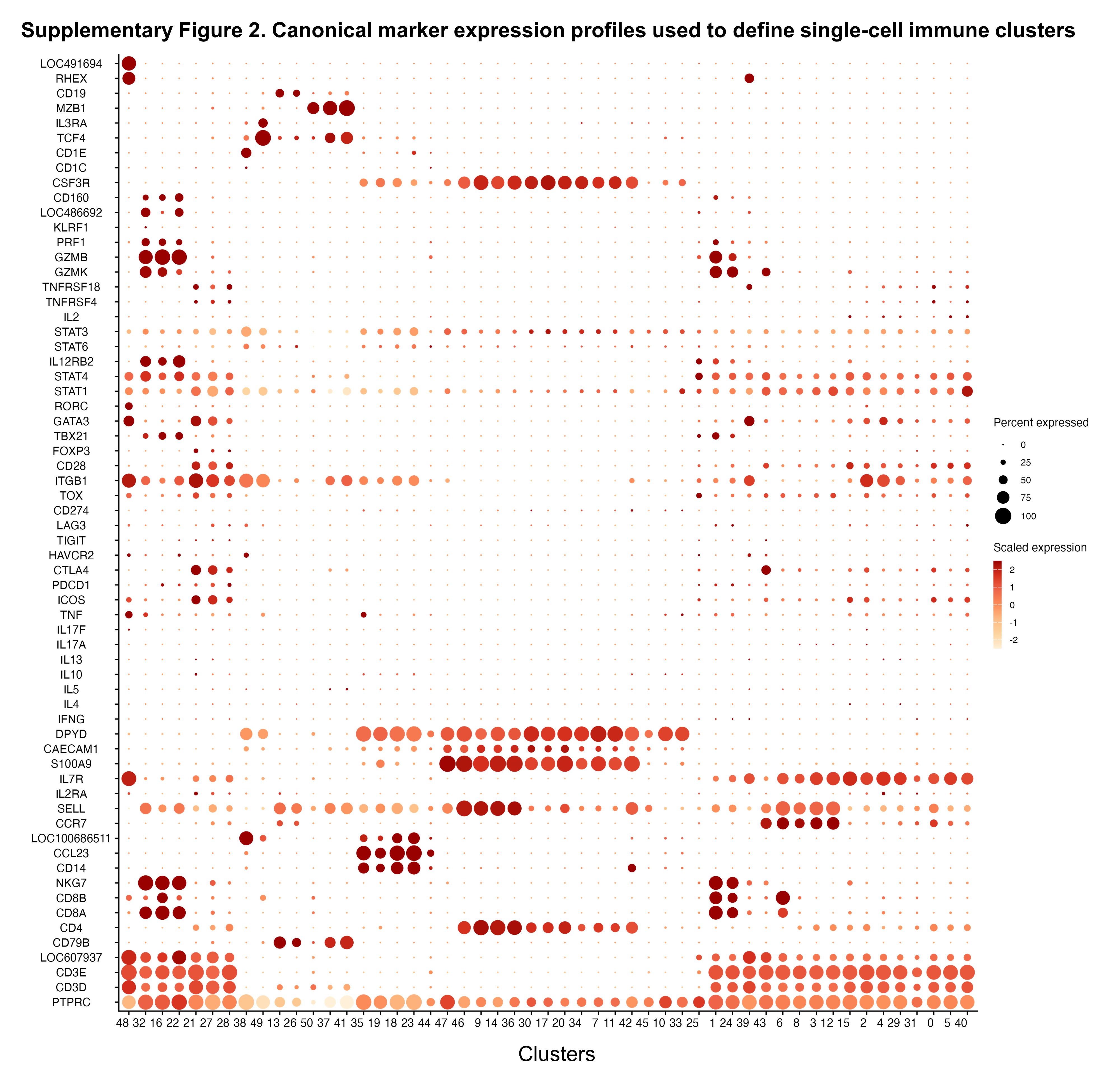

### Suppl_Figure_3.TIF

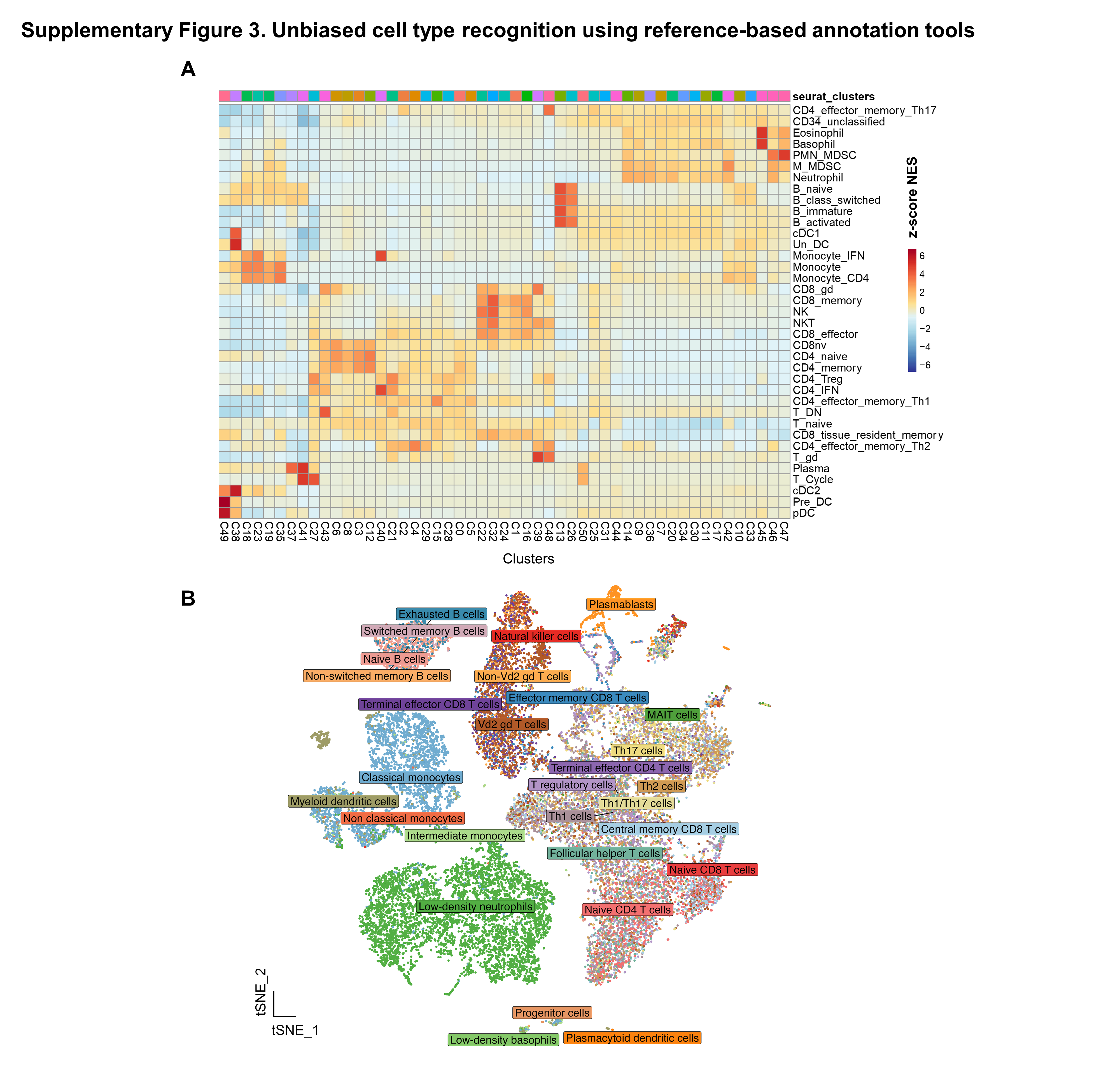

### Suppl_Figure_4.TIF

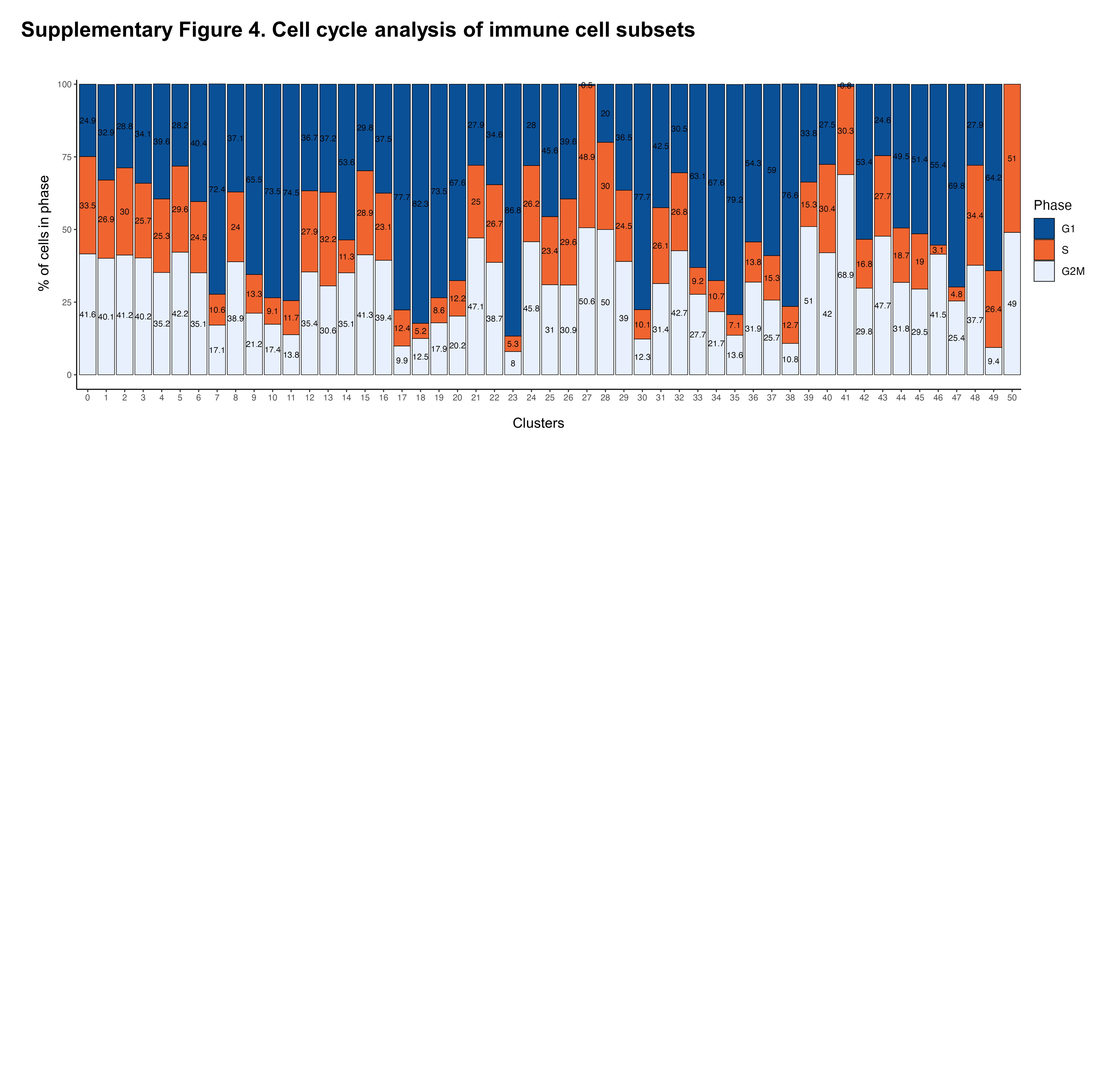

### Suppl_Figure_5.TIF

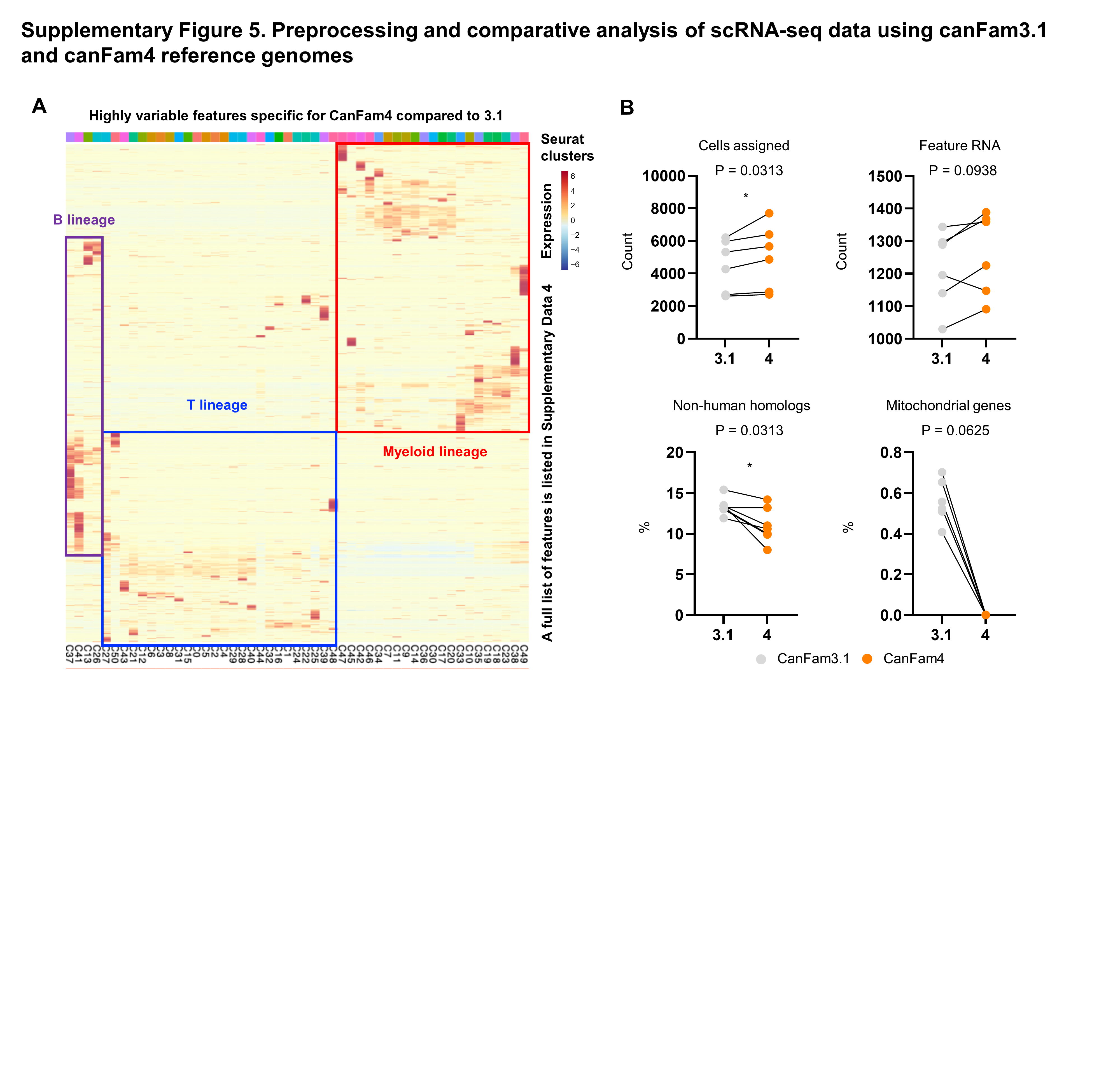

### Suppl_Figure_6.TIF

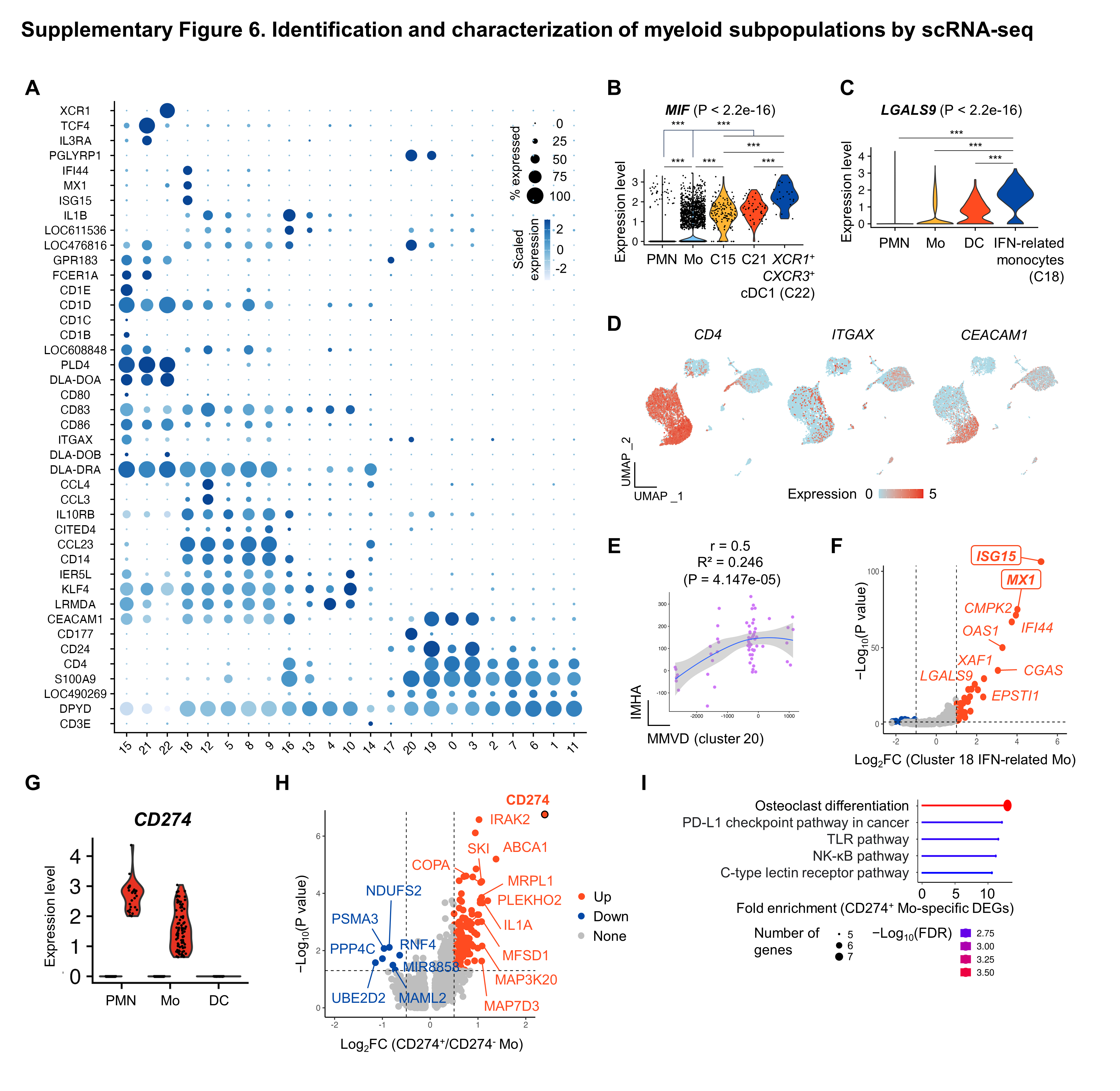

### Suppl_Figure_7.TIF

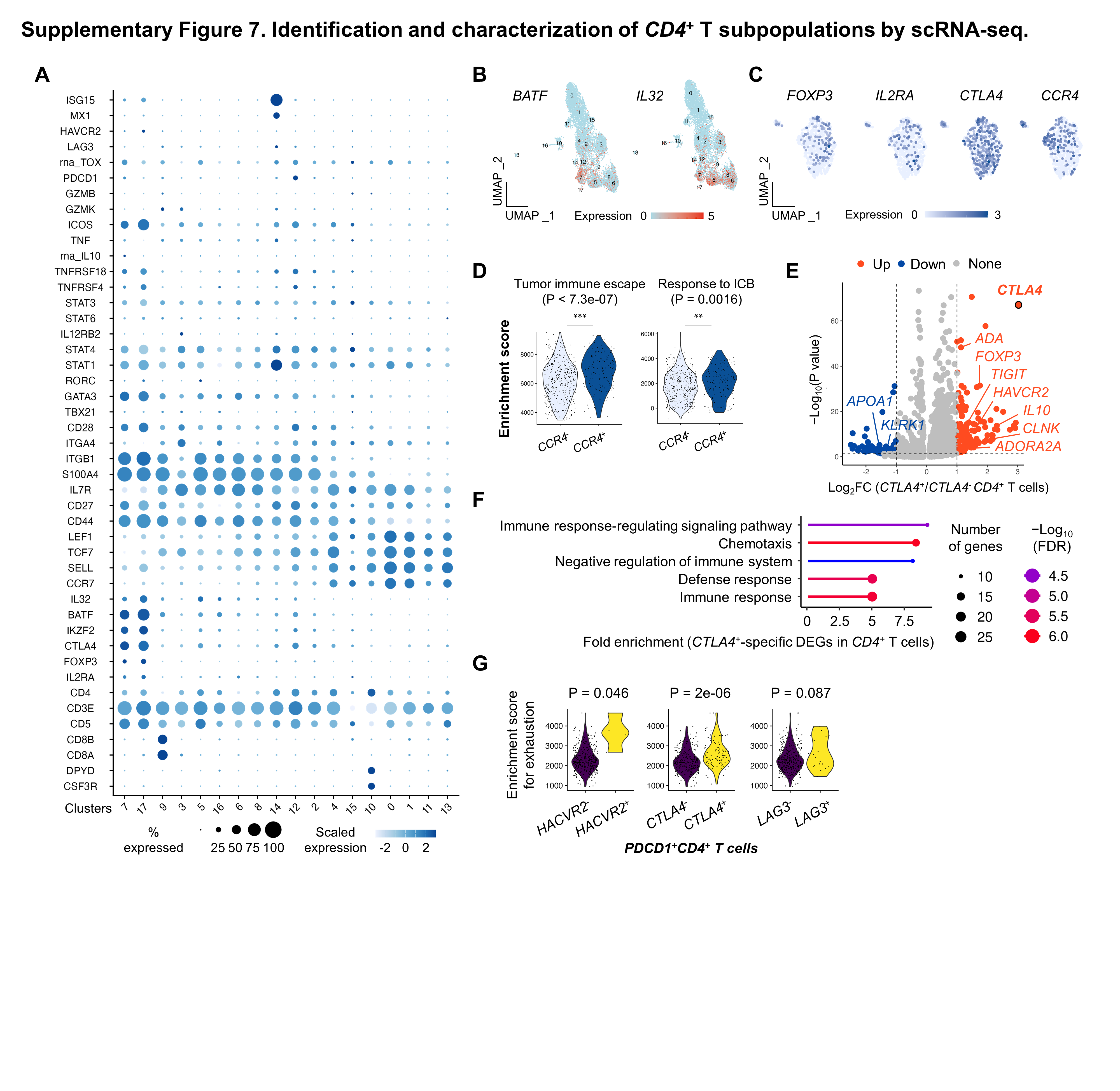

### Suppl_Figure_8.TIF

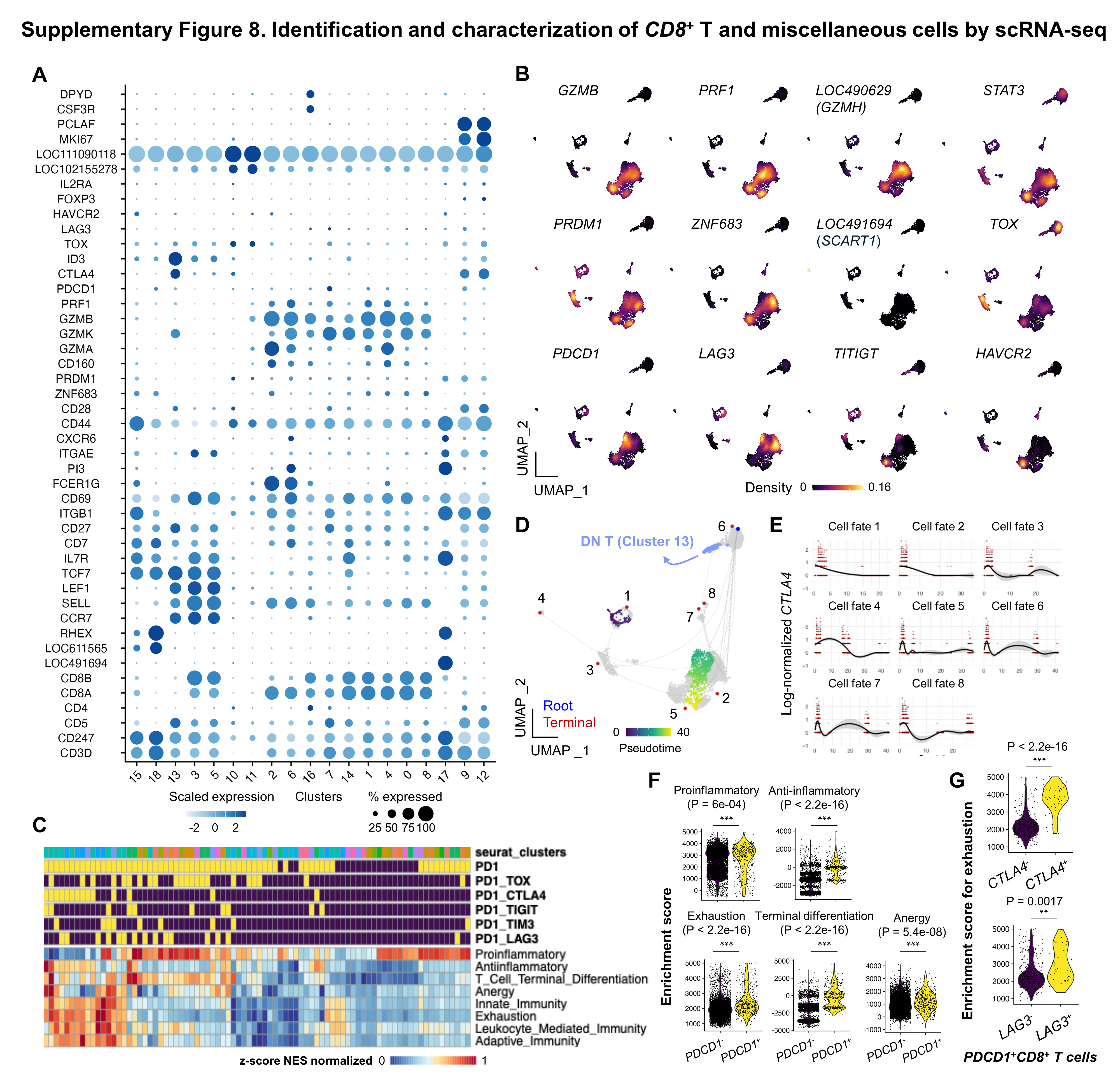

### Suppl_Figure_9.TIF

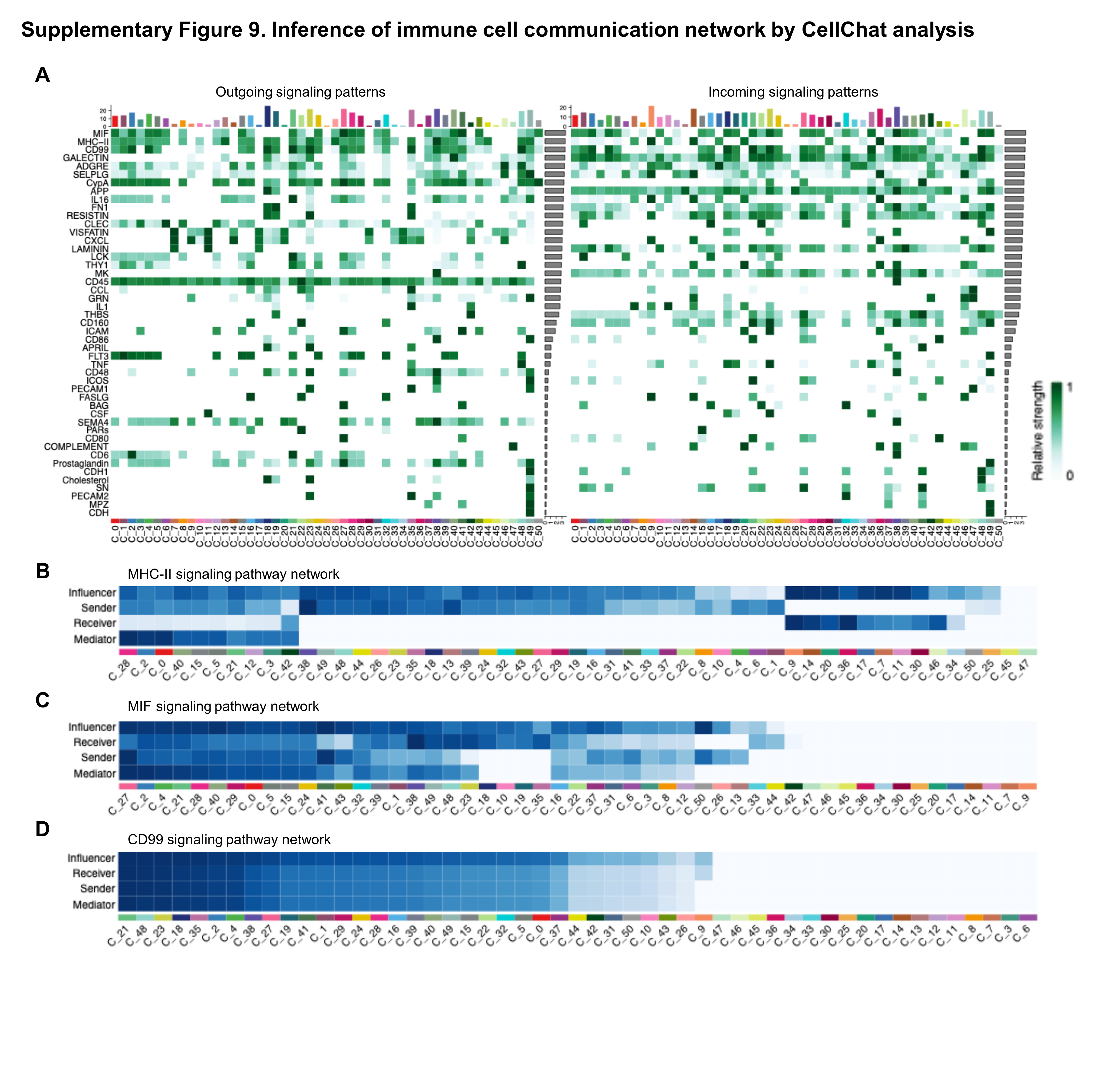

### Suppl_Figure_10.TIF

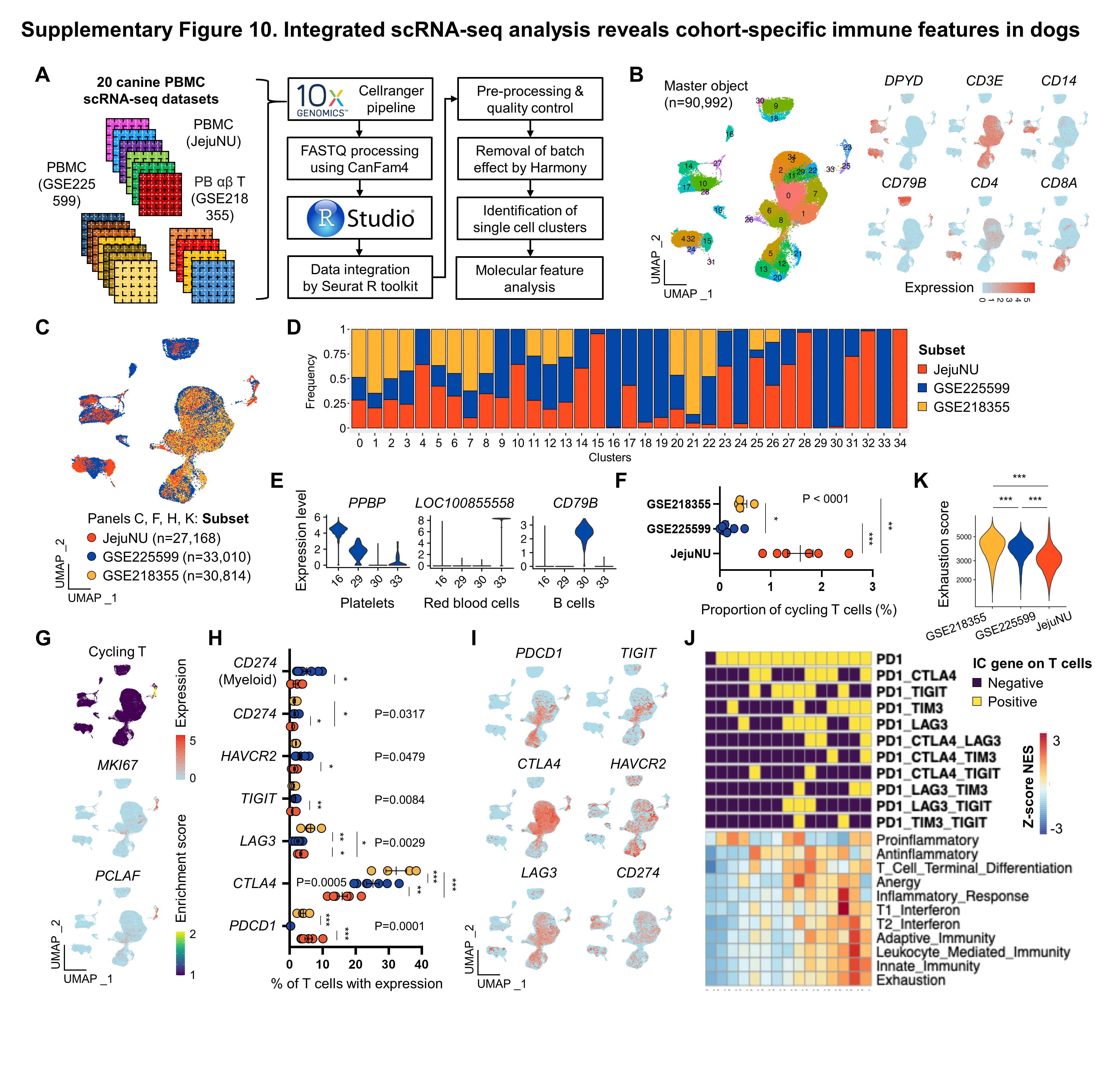
